# YPEL controls synapse development through p62-Nrf2 antioxidant pathway in *Drosophila* neuromuscular junction

**DOI:** 10.1101/2025.04.07.647271

**Authors:** Tianlu Wei, Ramin Lashanizadegan, Lily Liu, Monika Singh, Leanne Gayle L Mariano, Dennis Mathew, Jung Hwan Kim

## Abstract

The *Yippee-like* (*YPEL*) genes are highly conserved among all eukaryotic species, yet the molecular and cellular pathways that YPEL regulate are poorly understood. Human and animal studies suggest that YPEL is involved in nervous system functions. Here we report that YPEL is necessary for synapse development in neuromuscular junction and motor functions in *Drosophila*. YPEL interacts with the *Drosophila* p62, Refractory to sigma P, which forms cytoplasmic proteinaceous bodies for selective autophagy and signaling. YPEL overexpression decreased p62 bodies, while increased p62 bodies were observed in *YPEL* mutant neurons. Suppressing p62 bodies by reducing *p62* gene dosage significantly alleviated both synaptic and locomotion defects in *YPEL* mutants, suggesting that YPEL acts through p62 for synapse development. On the other hand, suppressing p62 bodies via autophagy did not restore synapse development in *YPEL* mutants. Interestingly, reduced levels of reactive oxygen species were found in *YPEL* mutants, which is consistent with the role of p62 in promoting the nuclear factor erythroid 2-related factor 2 (Nrf2) antioxidant pathway. Overexpressing *Drosophila* Nrf2, *Cap ‘n’ collar* (Cnc), phenocopied the synaptic and locomotion deficits in *YPEL* mutants. Importantly, both synaptic and locomotion defects were completely rescued by knocking down *Cnc* in *YPEL* mutant motor neurons. Taken together, our study demonstrates that YPEL negatively controls p62 - Nrf2 antioxidant pathway for neuromuscular synapse development and locomotion.

## Introduction

The *Yippee-like* (YPEL) gene family is highly conserved across a wide range of eukaryotic species [1], which suggests their fundamental cellular functions. There are five *YPEL* homologs in mice and humans based on amino acid sequence homology. YPEL1 to YPEL4 share very high sequence homology, while YPEL5 constitutes a distinct isoform. Studies suggested that YPEL proteins are involved in diverse functions ranging from cell senescence to tissue development. Mutations of *YPEL* genes caused defective fungus development [2]; YPEL1 is involved in fibroblast morphology and associated with Craniofacial Microsomia 1 and Branchiootic Syndrome [3, 4];YPEL2 and YPEL3 in cell senescence [5–8];YPEL4 in cell proliferation and red blood cell functions [9, 10]; YPEL5 in liver development in zebrafish [11] and cell cycle progression [12]. However, no clear mechanisms are known for these functions.

Studies suggest that YPEL3 functions in the nervous system as well. Various microdeletion and duplications in human chromosome 16p11.2, where *YPEL3* is located, are associated with multiple brain disorders including autism, schizophrenia, and a group of movement disorders [13–19]. A recent human case study identified a de novo frameshift mutation in *YPEL3* that is associated with various neurological symptoms including hypotonia and low muscle tone [20]. Although defective glial development and reduced synaptic transmission were described in zebrafish [20] and fly *YPEL3* mutants [21] respectively, the molecular and cellular mechanisms by which YPEL3 regulates nervous system development are largely unknown.

Sequestosome1, or p62/SQSTM1 is a widely expressed ubiquitin-binding protein that is found in most cytoplasmic inclusions in protein aggregation diseases including Alzheimer’s and Parkinson’s diseases [22]. Diverse missense mutations in *p62* are associated with amyotrophic lateral sclerosis [23]. On the other hand, *p62*-loss is associated with early onset neurodegeneration [24, 25]. Collectively, these suggest an important role of p62 in the nervous system and in the pathogenesis of neurodegenerative disorders. p62, through self-oligomerization, forms cytoplasmic proteaceous bodies, or p62 bodies that recruit ubiquitinated protein aggregate for autophagic degradation [26]. p62 bodies are also known to act as a signaling platform and regulate diverse signaling pathways including cellular antioxidant response via nuclear factor erythroid 2-related factor 2 (Nrf2) [27]. Through these mechanisms, p62 is involved in clearing cellular toxic aggregates and mitigates oxidative stress. Both autophagy and cellular redox are involved in neuronal functions [28, 29]. However, whether p62 is directly involved in synapse development is not known.

Here, we demonstrate that *dYPEL* - the sole ortholog of *YPEL1* to *YPEL4* in *Drosophila* - is essential for synapse development and locomotor activity in *Drosophila* larval neuromuscular junction. We found that dYPEL interacts with the *Drosophila* p62 and suppresses p62 bodies, hence an increase in p62 bodies upon *dYPEL*-loss, which was causative for the synaptic and locomotor deficits in *dYPEL* mutants. Our further analysis demonstrated that dYPEL negatively controls the cellular antioxidant pathway of p62, rather than autophagy. Heightened Nrf2 antioxidant response phenocopied the synaptic and locomotor deficits seen in *dYPEL* mutants. Conversely, inhibiting Nrf2 activity was sufficient to restore both synaptic and locomotor deficits by *dYPEL* mutations. Thus, our study defines a previously unknown role of YPEL in synapse development and YPEL as a negative regulator of the p62-antioxidant pathway.

## Result

### dYPEL is required for synapse development in *Drosophila* NMJ and locomotor activity

A mutation in *YPEL3* was identified in a patient with a rare neurological disorder characterized by several neurological symptoms, including areflexia and hypotonia [20]. Our group previously reported reduced synaptic transmission in sensory neurons in *Drosophila YPEL* (*dYPEL*) mutants [21]. *dYPEL* represents a sole ortholog for YPEL1 to YPEL4 in *Drosophila*. Although this provided evidence for a potential synaptic role of YPEL, how YPEL controls synaptic transmission and whether YPEL is required for synapse development is not known. Since both areflexia and hypotonia may be associated with muscle weakness, we examined if dYPEL is involved in synapse development of larval neuromuscular junction (NMJ). Two independent *dYPEL* mutant alleles were analyzed: *dYPEL^T1-8^*, a frameshift mutation causing a premature stop codon, which harbors similar molecular lesions that were found in a human patient, and *dYPEL^KO^*, in which the entire *dYPEL* open reading frame was deleted, hence represents molecular null allele [21]. An anti-Horseradish-peroxidase (HRP) antibody and an anti-Bruchpilot (BRP) antibody were used to visualize the NMJ and presynaptic active zones, respectively. The type Ib bouton was imaged from muscle 4. The analysis revealed dramatic defects in NMJ development. The number of presynaptic active zones was reduced by 40% and 33% in *dYPEL^T1-8^* and *dYPEL^KO^*, respectively. Similarly, the NMJ area was reduced by 28% and 22% in *dYPEL^T1-8^* and *dYPEL^KO^*, respectively, as compared to controls (**Figure 1A**).

**Figure 1.**
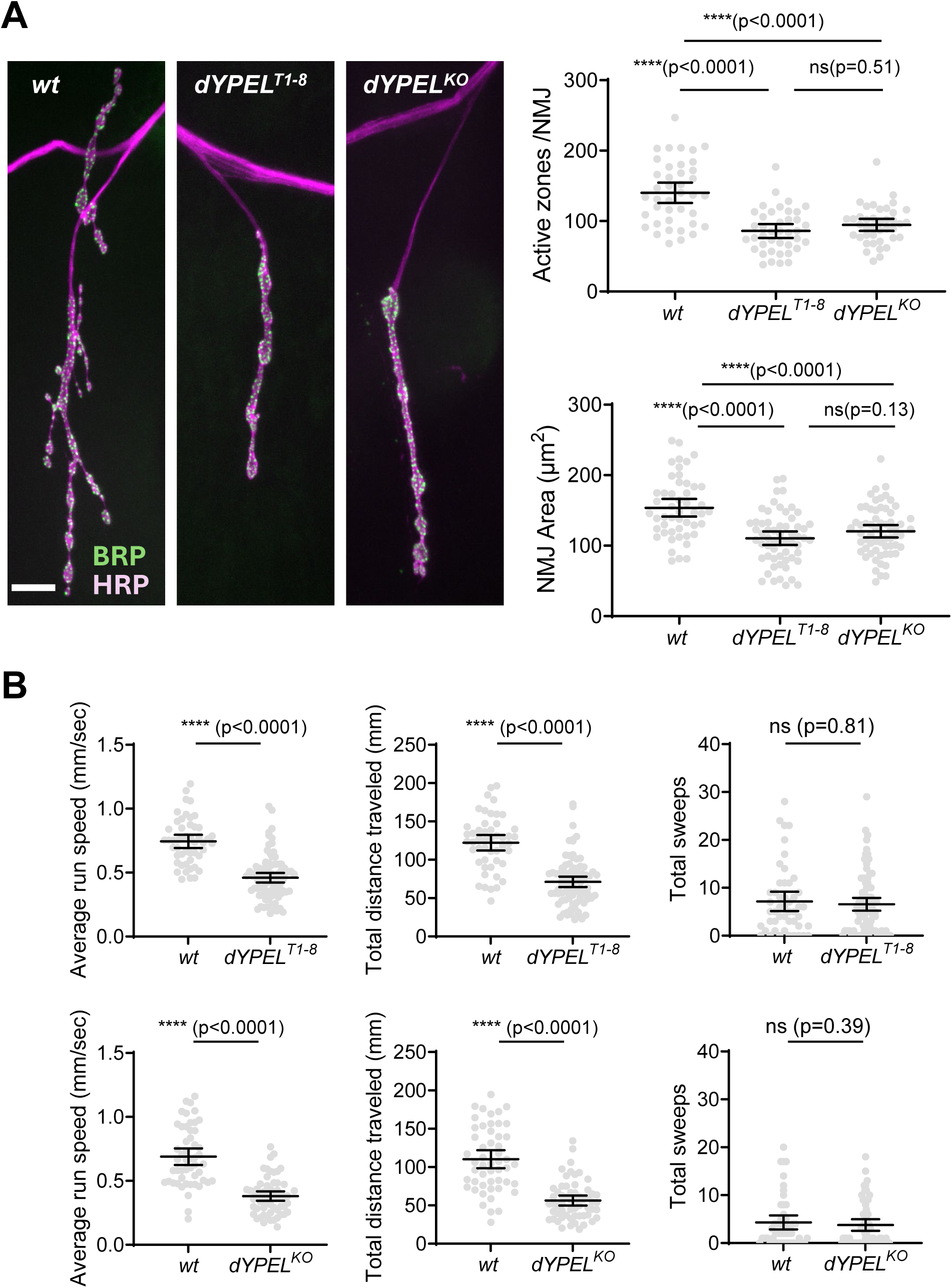
*dYPEL* is necessary for neuromuscular synaptic development and locomotor activity in *Drosophila* larva. **(A)** NMJ analysis of *dYPEL* mutants. Left: Representative NMJ images from *w^1118^*, *dYPEL^T1-8^* and *dYPEL^KO^*, images were obtained from the type Ib boutons on muscle 4 from third instar male larvae. Anti-HRP staining was used to label NMJ (magenta). Anti-BRP staining was used to label presynaptic active zones (green). Scale bar = 10 μm. Top right: *dYPEL* mutations caused a reduction in active zones. The total number of BRP-positive puncta, or “Active zone” was quantitated from the type Ib NMJ images and expressed as mean ± 95% CI. The genotypes and sample numbers were *w^1118^* (n = 40), *dYPEL^T1-8^* (n = 40), and *dYPEL^KO^* (n = 40). One-way ANOVA (F(2,117) =27.67) followed by post hoc Turkey’s multiple comparison test. Bottom right: *dYPEL* mutations caused smaller NMJ sizes. The total NMJ area was calculated from anti-HRP staining. and expressed as mean ± 95% CI. An automatic annotation was used to define the NMJ area (see Materials and Method). The genotypes and sample numbers were *w^1118^* (n = 49), *dYPEL^T1-8^* (n = 59), and *dYPEL^KO^* (n = 62). One-way ANOVA (F(2,167) =18.64) followed by post hoc Turkey’s multiple comparison test. **(B)** *dYPEL* mutations caused reduced locomotor activity. Locomotor behavior analysis of *dYPEL^T1-8^* and *dYPEL^KO^*. The third instar larvae were placed in a crawling behavior arena. Individual larva crawling was traced and quantified using a custom code to obtain the average run speed, the total distance traveled, and total number of sweeps. Data was presented as mean ± 95% CI. The genotypes and sample sizes for the top row were *w^1118^* (n = 49), *dYPEL^T1-8^* (n = 84). The two-tailed Mann-Whitney test was used (U = 502, p<0.0001 for the average run speed; U=567, p<0.0001 for the total distance traveled; U=2007, p=0.81 for the total sweeps). The genotypes and samples for the bottom row were *w^1118^* (n = 51), *dYPEL^KO^* (n = 59). Data was presented as mean ± 95% CI. The two-tailed Mann-Whitney test was used (U = 349, p<0.0001 for the average run speed; U=383, p<0.0001 for the total distance traveled; U=1364, p=0.39 for the total sweeps).

To test if defective NMJ development in *dYPEL* mutants leads to locomotor defects, we measured the larva crawling behavior. The behavior was measured under complete darkness and essentially in the absence of sensory stimulation. Individual larva crawling was traced and analyzed as previously described [30]. The result showed a dramatic reduction in larva locomotion by *dYPEL* mutations. The *dYPEL^T1-8^* mutants exhibited a significant 38% reduction in average run speed and a 42% decrease in the total distance travelled compared to controls (**Figure 1B**). Similarly, *dYPEL^KO^* larvae displayed a significant 44% reduction in the average run speed and a 49% reduction in the total distance travelled compared to controls (**Figure 1B**). In contrast, the total number of head sweeps was comparable between controls and *dYPEL* mutants (**Figure 1B**). The head sweeping behavior is associated with food searching, exploration, defensive actions and social interactions, which reflect the internal status of larva [30]. Together, these demonstrated that dYPEL is necessary for synaptic development in the NMJ and locomotion.

### dYPEL regulates Ref(2)P

To investigate the cellular and molecular mechanisms by which dYPEL regulates synapse development and locomotion, we examined potential dYPEL binding proteins. A proteome-wide interaction analysis has identified Refractory to Sigma P, or Ref(2)P, as a potential dYPEL-interacting protein in *Drosophila* [31]. Ref(2)P is the *Drosophila* ortholog of mammalian p62/SQSTM1.

To confirm the interaction between dYPEL and Ref(2)P, we performed co-immunoprecipitation (Co-IP) using *Drosophila* Schneider 2 cells (S2). Myc-tagged dYPEL and GFP-tagged Ref(2)P were co-expressed in S2 cells. Ref(2)P::GFP was immunoprecipitated using an anti-GFP nanobody, and the presence of bound dYPEL::myc was detected by Western blot analysis. Cells transfected with dYPEL::myc and an empty plasmid DNA served as a negative control. We observed robust pull-down of dYPEL::myc by Ref(2)P::GFP (**Figure 2A**). In contrast, the GFP nanobody did not pull down dYPEL::myc in the absence of Ref(2)P::GFP. The expression levels of dYPEL::myc were not different in both groups. This confirms that dYPEL physically interacts with Ref(2)P in *Drosophila*.

**Figure 2.**
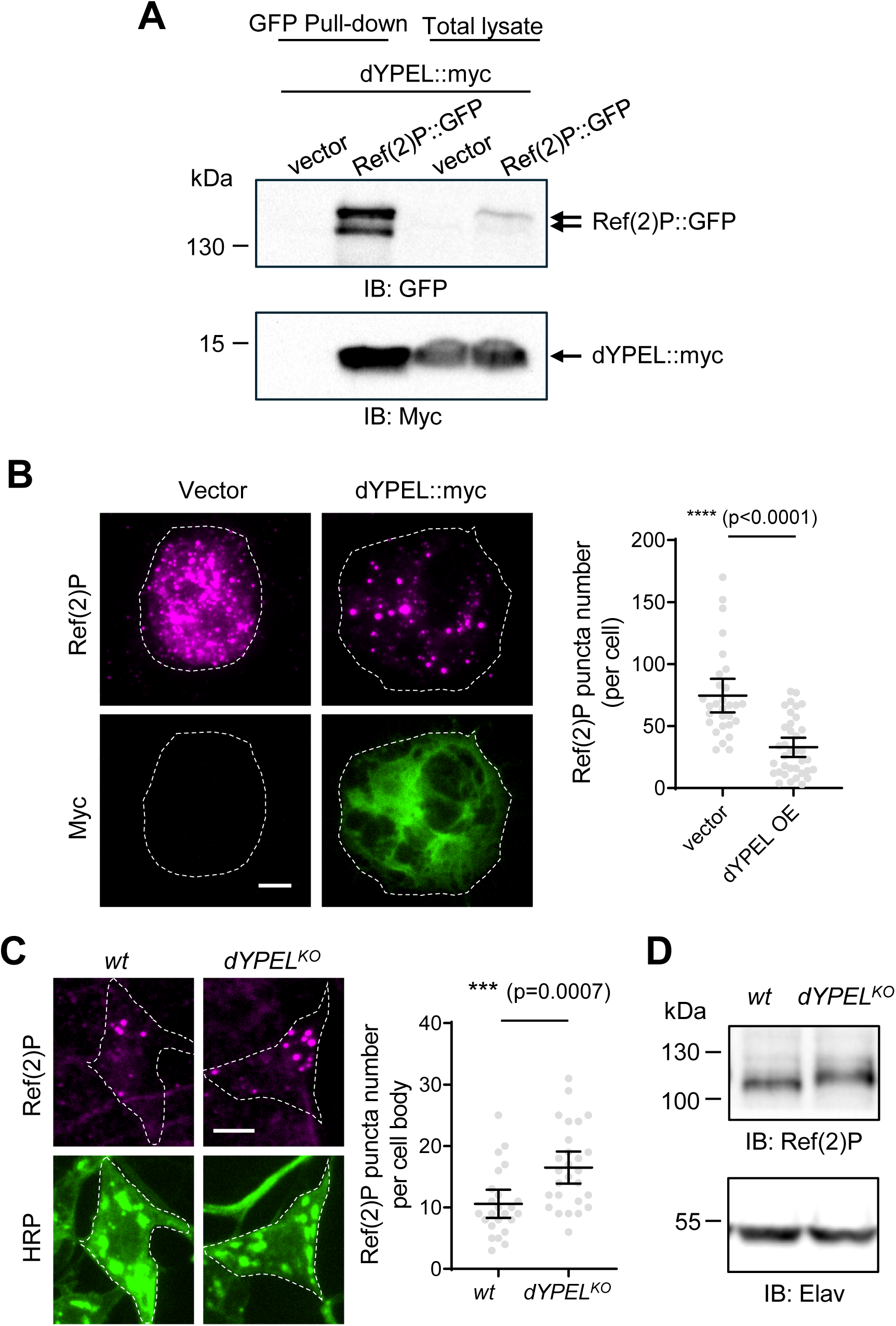
*dYPEL* regulates Ref(2)P bodies. **(A)** *dYPEL* physically interacts with Ref(2)P. dYPEL::myc and Ref(2)P::GFP were co-transfected into S2 cells. Ref(2)P::GFP was pulled down using anti-GFP nanobody (GFP Pull-down). The input (Total lysate) was loaded as a comparison. An empty plasmid (vector) served as a negative control. The Western blotting was performed using anti-GFP and anti-Myc antibodies to detect Ref(2)P::GFP and dYPEL::myc respectively. Similar results were obtained in two other independent repeats. **(B)** *dYPEL* reduces the number of Ref(2)P bodies in S2 cells. Representative images of Ref(2)P puncta in S2 cells. Either dYPEL::myc or an empty plasmid (vector) was transfected into S2 cells. An anti-Ref(2)P antibody was used to visualize endogenous Ref(2)P proteins (magenta). An anti-Myc antibody was used to identify dYPEL::myc expressing cells (green). The Ref(2)P-positive puncta were quantitated and presented as mean ± 95% CI. The two-tailed Mann-Whitney test was used (U = 154.5, p<0.0001). Sample numbers were n = 29 for vector and n = 36 for dYPEL::myc. Scale bar = 10μm. **(C)** *dYPEL* mutations increased Ref(2)P bodies in C4da neurons. Representative images of Ref(2)P staining in C4da neurons from wild type control (*wt*; *w^1118^*) and *dYPEL^KO^*. An anti-Ref(2)P antibody was used to visualize Ref(2)P bodies (magenta). C4da neurons were identified using HRP staining (green). Scale bar = 10 μm. The Ref(2)P-positive puncta were quantitated and presented as mean ± 95% CI. The two-tailed Mann-Whitney test was used (U = 148.5, p=0.0007). Sample numbers were *dYPEL^KO^* (n=27) and *w^1118^* (n=24). **(D)** *dYPEL* mutations do not affect Ref(2)P protein levels. The total lysates from the third-instar larvae brain from wt (*w^1118^*) and *dYPEL^KO^* were subjected to SDS-PAGE and Western blot analysis. An anti-Ref(2)P antibody was used to detect endogenous Ref(2)P protein levels. Anti-Elav staining was used as a loading control. Similar results were obtained in three other independent repeats.

p62/Ref(2)P is known to form cytoplasmic proteinaceous bodies. A recent study reported that YPEL2 and p62 colocalize in stress granules under cellular stress [32]. To determine whether dYPEL interacts with Ref(2)P in these proteinaceous bodies, we performed colocalization analysis in S2 cells. Distinct cytoplasmic puncta structures were observed by Ref(2)P antibody staining in S2 cells. On the other hand, dYPEL::myc showed diffuse cytoplasmic staining that often excludes Ref(2)P puncta (**Figure 2B**), suggesting that dYPEL proteins interact with soluble Ref(2)P proteins. Interestingly, we observed a dramatic 56% reduction in Ref(2)P puncta number in dYPEL::myc expressing cells as compared to the empty plasmid transfected control groups. These suggest that YPEL negatively controls p62/Ref(2)P bodies.

To assess whether dYPEL regulates Ref(2)P bodies in neurons in vivo, we immunostained wild-type and *dYPEL^KO^* larvae with an anti-Ref(2)P antibody. Extensive non-specific staining by the anti-Ref(2)P antibody in the ventral nerve cord precluded us from examining Ref(2)P bodies in the motor neurons. Instead, we analyzed a sensory neuron type (Class IV dendritic arborization neuron/C4da neuron), where Ref(2)P puncta were readily observed in their cell bodies. A significant 56% increase in Ref(2)P puncta number was observed in *dYPEL^KO^* C4da neurons (**Figure 2C**), indicating that dYPEL negatively regulates Ref(2)P puncta formation in vivo. dYPEL may regulate Ref(2)P bodies by repressing Ref(2)P protein levels. Western blot analysis on the larval brain lysates did not show changes in Ref(2)P protein levels by *dYPEL^KO^* (**Figure 2D**). We concluded that dYPEL regulates Ref (2)P bodies without affecting Ref(2)P proteins levels.

### dYPEL regulates synapse development and locomotor activity through Ref(2)P

The direct involvement of p62/Ref(2)P in synapse development is not known. p62/Ref(2)P is involved in selective autophagy as well as in various signaling events [27]. Moreover, mutations in *p62* or abnormal accumulation of p62 bodies were associated with neurodegenerative conditions [22–25]. Thus, we wondered whether increased Ref(2)P bodies are causative for the deficits in synapse development and locomotion by *dYPEL* mutations. To answer the question, we attempted to normalize Ref(2)P puncta number by reducing *Ref(2)P* gene dosage. *Ref(2)P^OD2^* is a loss-of-function allele that lacks the self-oligomerization domain of Ref(2)P [33], thus presumably preventing Ref(2)P body formation. Introducing one copy of *Ref(2)P^OD2^* in *dYPEL^KO^* C4da neurons (*dYPEL^KO^, Ref(2)P^OD2^/+*) reduced Ref(2)P puncta number similar to that from a wild-type control while *Ref(2)P^OD2^* heterozygous itself did not affect Ref(2)P puncta number in wild-type C4da neurons (**Figure 3A**). Next, we measured presynaptic active zones in a wild type, *dYPEL^KO^*, *Ref(2)P^OD2^/+*, and *dYPEL^KO^, Ref(2)P^OD2^/+* NMJ using anti-BRP antibody. While introducing one copy of *Ref(2)P^OD2^* did not significantly alter synaptic active zones, it restored active zone numbers from a 62% to 82% of the control (*dYPEL^KO^* and *dYPEL^KO^, Ref(2)P^OD2^/+*) (**Figure 3B**).

**Figure 3.**
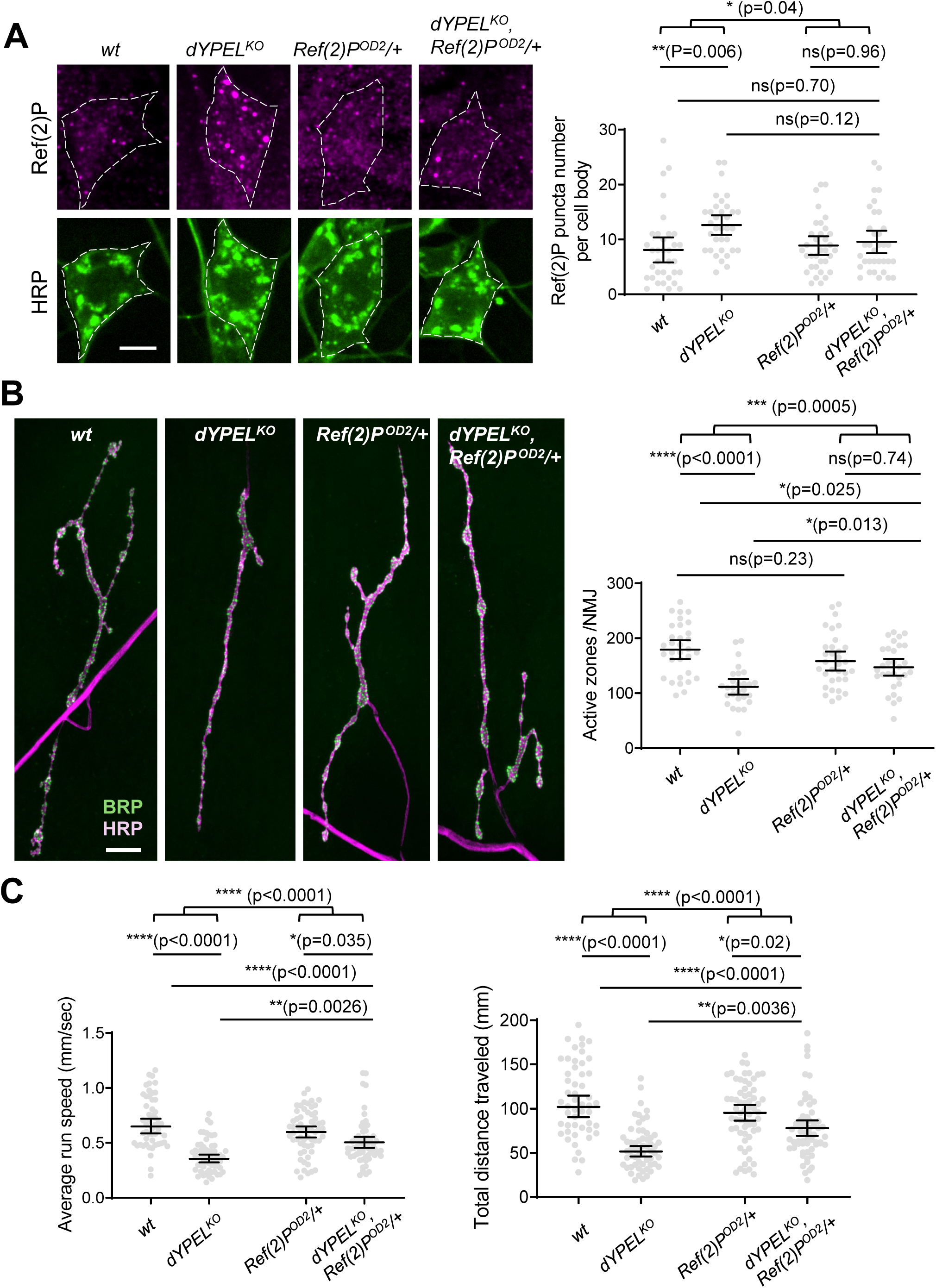
*dYPEL* regulates synapse development and locomotor activity through Ref(2)P. **(A)** Ref(2)P bodies in *dYPEL* mutants were normalized by reducing *Ref(2)P* gene dosage. An anti-Ref(2)P antibody was used to visualize Ref(2)P bodies in *wt* (*w^1118^*), *dYPEL^KO^*, *Ref(2)P^OD2^/+*, and *dYPEL^KO^*, *Ref(2)P^OD2^/+* C4da neurons (magenta). HRP staining was used to identify C4da neurons (green). Scale bar = 10μm. Samples were obtained from third-instar male larvae. The Ref(2)P-positive puncta were quantitated and presented as mean ± 95% CI. The genotypes and sample numbers were: *w^1118^* (n = 34); *dYPEL^KO^*(n = 34); *Ref(2)P^OD2^*/+ (n = 34); *dYPEL^KO^*, *Ref(2)P^OD2^*/+ (n = 34). Two-way ANOVA ((F1,132) =3.99) followed by post hoc Turkey’s correction. **(B)** Reduced *Ref(2)P* gene dosage alleviated synaptic defect in *dYPEL* mutant NMJ. An anti-BRP antibody was used to visualize presynaptic active zones in *wt* (*w^1118^*), *dYPEL^KO^*, *Ref(2)P^OD2^/+*, and *dYPEL^KO^*, *Ref(2)P^OD2^/+* NMJ (green). HRP staining was used to visualize the type Ib boutons (magenta). Samples were obtained from third-instar male larvae. Scale bar = 10 μm. The total number of BRP-positive puncta, or “Active zone” was quantitated from the type Ib NMJ images and expressed as mean ± 95% CI. The genotypes and sample numbers were *w^1118^* (n = 32); *dYPEL^KO^* (n = 28); *Ref(2)P^OD2^/+* (n = 33); *dYPEL^KO^*, *Ref(2)P^OD2^/+* (n = 30). Two-way ANOVA ((F1,119) =12.67) followed by post hoc Turkey’s correction. **(C)** Reduced *Ref(2)P* gene dosage alleviated locomotor defect in *dYPEL* mutants. The third instar male larvae were placed in a crawling behavior arena. Individual larva crawling was traced and quantified using a custom code to obtain the average run speed and the total distance traveled. Data was presented as mean ± 95% CI. The genotypes and sample numbers were *w^1118^* (n = 51); *dYPEL^KO^* (n = 59); *Ref(2)P^OD2^/+* (n = 62); *dYPEL^KO^*, Ref(2)P^OD2^/+ (n = 63). Average run speed: Two-way ANOVA (F(1,231) =18.13) followed by post hoc Turkey’s correction. Total distance travelled: Two-way ANOVA (F(1,231) =16.57) followed by post hoc Turkey’s correction.

We next measured larva locomotion in the presence of one copy of *Ref(2)P^OD2^*. The result showed that *Ref(2)P^OD2^/+* partially but significantly rescued the locomotor deficits induced by *dYPEL^KO^*. The average run speed was restored to 74% from 55% of wild-type and the distance travelled was restored to 70% from 51% by *Ref(2)P^OD2^/+* (**Figure 3C**). Introducing a copy of *Ref(2)P^OD3^*, another *Ref(2)P* loss-of-function allele led to a partial rescue of locomotor activity in *dYPEL^KO^* (**Figure 3-supplement**). Taken together, these results suggest that p62/Ref(2)P bodies are downstream of YPEL in synapse development and locomotor activity.

### dYPEL regulates NMJ development independent of autophagy

p62/Ref(2)P is a multifunctional protein involved in several cellular processes, particularly in the regulation of selective autophagy. The p62/Ref(2)P bodies sequester cytoplasmic aggregates in them and promote autophagosome formation [26]. Thus, p62/Ref(2)P bodies undergo autophagic degradation along with the autophagy cargos [26]. Thus, the increased number of Ref(2)P puncta seen in *dYPEL* mutants may be caused by impaired autophagy. To test if increased autophagy can normalize Ref(2)P puncta number as well as the NMJ synaptic defects in *dYPEL* mutants, we expressed Autophagy-related 1 (Atg1) in C4da neurons under a C4da-specific driver *ppk-Gal4* [34]. Atg1 is a key kinase that initiates autophagy [35]. Quantitative analysis of Ref(2)P puncta in C4da cell bodies revealed that Atg1 overexpression brought back the increased Ref(2)P puncta number observed in *dYPEL^KO^* to the wild-type levels (**Figure 4A**).

**Figure 4.**
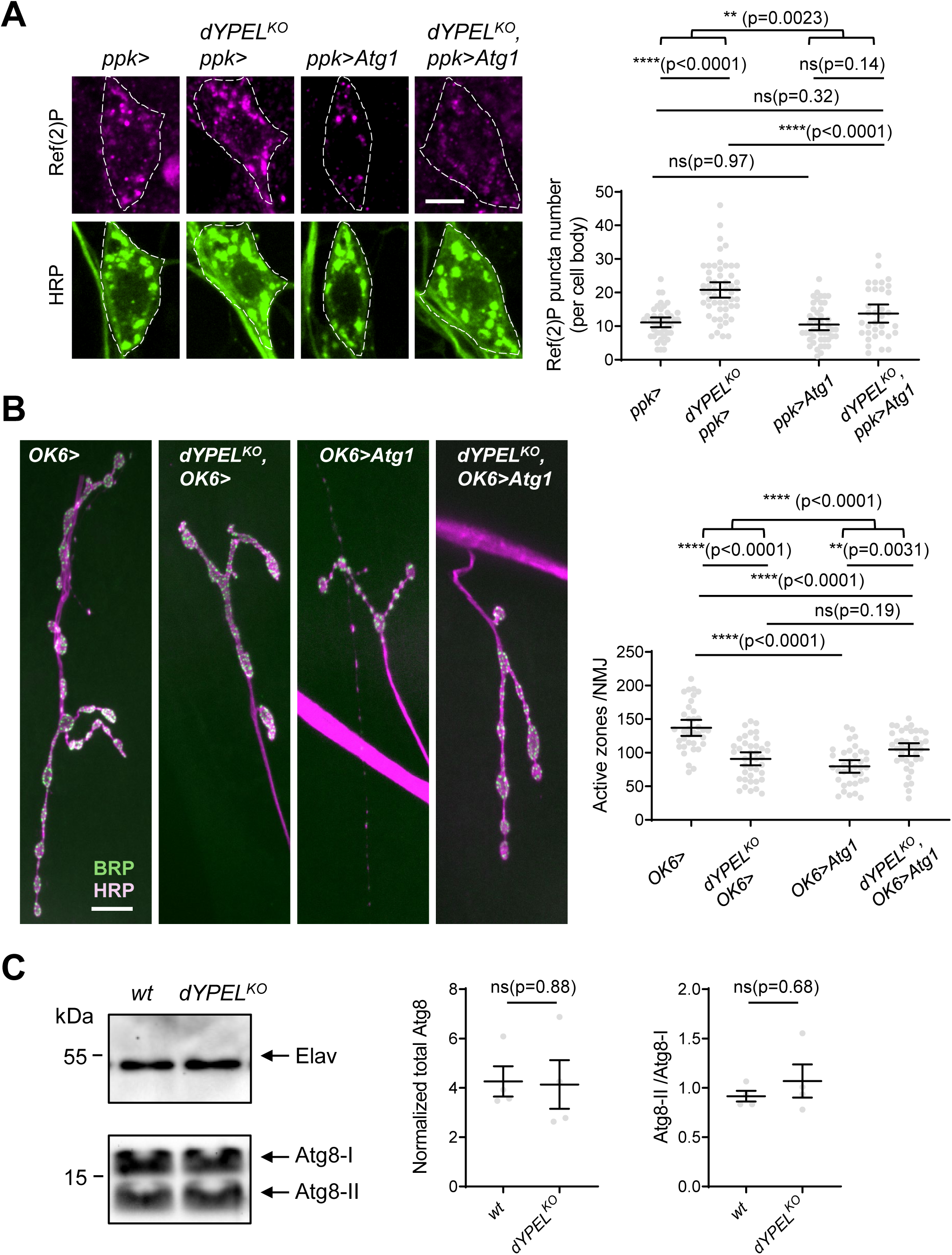
*dYPEL* regulates neuromuscular synapse development independently of autophagy. **(A)** Increased autophagy suppresses Ref(2)P body increase in *dYPEL* mutant C4da neurons. Atg1 was expressed in C4da neurons under the C4da-specific driver *ppk-Gal4.* Representative images of wild type control (*w^1118^, ppk>*), *dYPEL^KO^* (*dYPEL^KO^*, *ppk>*), Atg1 overexpression (*ppk>*Atg1), and Atg1 overexpression in *dYPEL^KO^* (*dYPEL^KO^, ppk>*Atg1) are shown. Ref(2)P puncta were labeled with an anti-Ref(2)P antibody (magenta). C4da neurons were identified with HRP staining (green). Samples were obtained from third-instar male larvae. Scale bar = 10μm. The Ref(2)P-positive puncta were quantitated and presented as mean ± 95% CI. The genotypes and sample numbers were *ppk>* (n = 43); *dYPEL^KO^*, *ppk>* (n = 53); *ppk>*Atg1 (n = 46); *dYPEL^KO^, ppk>*Atg1 (n = 34). Two-way ANOVA (F(1, 172) = 9.549) followed by post hoc Turkey’s correction. **(B)** Altered synapse development by autophagy in wild type and *dYPEL^KO^* NMJ. Atg1 was expressed in motor neurons using the *OK6-Gal4* driver. An anti-BRP antibody was used to visualize presynaptic active zones (green). HRP staining was used to visualize the type Ib boutons (magenta). Representative NMJ images of wild type control (*w^1118^, OK6>*), *dYPEL^KO^* (*dYPEL^KO^*, *OK6>*), Atg1 overexpression (*OK6>Atg1*), and Atg1 overexpression in *dYPEL^KO^* (*dYPEL^KO^, OK6>Atg1*). Scale bar = 10μm. Samples were obtained from third-instar male. The total number of BRP-positive puncta, or “Active zone” was quantitated from the type Ib NMJ images and expressed as mean ± 95% CI. The genotypes and sample numbers were: *OK6>* (n = 36); *dYPEL^KO^,OK6>* (n = 40); *OK6>Atg1* (n = 36); *dYPEL^KO^*, *OK6>Atg1* (n = 39). Two-way ANOVA (F(1, 122) = 30.39) followed by post hoc Turkey’s correction. **(C)** dYPEL does not affect autophagy. The lysates from wild type control (*wt*, *w^1118^*) and *dYPEL^KO^* adult fly heads were subjected to Western blot analysis using an anti-Atg8 antibody. Elav blot was used as a loading control. The total intensity of Atg8 blot (Atg8-I + Atg8-II) was normalized by Elav total intensity to express the total Atg8 levels while the total intensity of Atg8-II was divided by that of Atg8-I to calculate Atg8-II/Atg8-I ratio. Data was presented as mean ± SEM. The two tailed Mann-Whitney test were used (U = 7, p = 0.88 for Total Atg8 levels; U = 6, p = 0.68 for Atg8-II/Atg8-I ratio). Sample numbers were 4 for both *w*t and *dYPEL^KO^*.

Next, we expressed Atg1 in motor neurons using *OK6-Gal4 36* [36] in wild-type and *dYPEL^KO^* larvae and analyzed the NMJ. Atg1 overexpression led to a dramatic reduction in active zones and smaller NMJ sizes — a 42% reduction in active zones and a 49% reduction in the NMJ area (**Figure 4B**), which was an even bigger reduction than in *dYPEL^KO^*. On the other hand, the NMJs from Atg1 overexpressing *dYPEL^KO^* motor neurons showed more active zones than Atg1 overexpression or *dYPEL^KO^* alone — yet still a dramatic reduction as compared to the wild-type control. These analyses suggest that there is a complex relationship between YPEL and autophagy pathway for NMJ synapse development.

To gain a better insight into the role of autophagy in YPEL function, we measured lipidation of the microtubule-associated protein 1 light chanin3 (LC3). LC3 lipidation is an essential process for phagophore formation [35]. Atg8a is the *Drosophila* LC3. The ratio between Atg8a-I, the cytoplasmic form and Atg8a-II, the lipidated form is commonly used as a maker for autophagic activity [37]. Adult fly heads from wild-type control and *dYPEL^KO^* were collected and processed for Western blot analysis. An anti-Atg8 antibody was used to detect endogenous Atg8-I and Atg8-II protein levels. Elav staining was used as the loading control. Western blot quantification revealed that both the total Atg8 expression levels and the Atg8-II/Atg8-I ratio were not significantly different between the control and *dYPEL^KO^* flies (**Figure 4C**), which indicates that dYPEL does not regulate autophagy.

Together, our data supports a model wherein dYPEL regulates NMJ synapse development through a pathway independent of autophagy.

### dYPEL regulates cellular antioxidant pathways

Aside from the functions in autophagy, p62/Ref(2)P is involved in various signaling events [27]. p62/Ref(2)P accumulates under stress conditions and enhances cellular antioxidant response as a protective mechanism through Kelch-like ECH-associated protein 1 (Keap1) - Nuclear factor erythroid 2-related factor 2 (Nrf2) pathway [27]. Neural activity affects synapse development by increasing reactive oxygen species (ROS) [29]. Some neuronal cell types suppress Nrf2 expression as they differentiate into mature neurons [38]. These suggest that certain ROS levels are necessary for neuronal development. These raise a possibility that YPEL regulates synapse development by altering cellular redox status through p62 - Nrf2 pathway. We employed a redox-sensitive probe roGFP to measure ROS levels in *dYPEL* mutants. The roGFP contains two cysteine residues into the GFP beta-barrel structure, allowing it to change fluorescence excitation spectra based on its oxidation state [39]. We expressed the mitochondrially targeted hydrogen peroxide sensor, mito-roGFP2-Orp1 [39] in wild-type and *dYPEL^KO^* larval motor neurons using *OK6-Gal4*. Green fluorescence confocal microscope images were sequentially acquired from the motor neuron cell bodies using 405 nm and 488 nm excitation. The relative hydrogen peroxide levels were calculated from the ratiometric analysis. The result showed that *dYPEL* mutations significantly reduced mitochondrial hydrogen peroxide levels (**Figure 5A**), which suggests that dYPEL negatively regulates cellular antioxidant pathway of Nrf2 by suppressing Ref(2)P bodies. This also suggests that *YPEL*-loss leads to overactive Nrf2 pathway.

**Figure 5.**
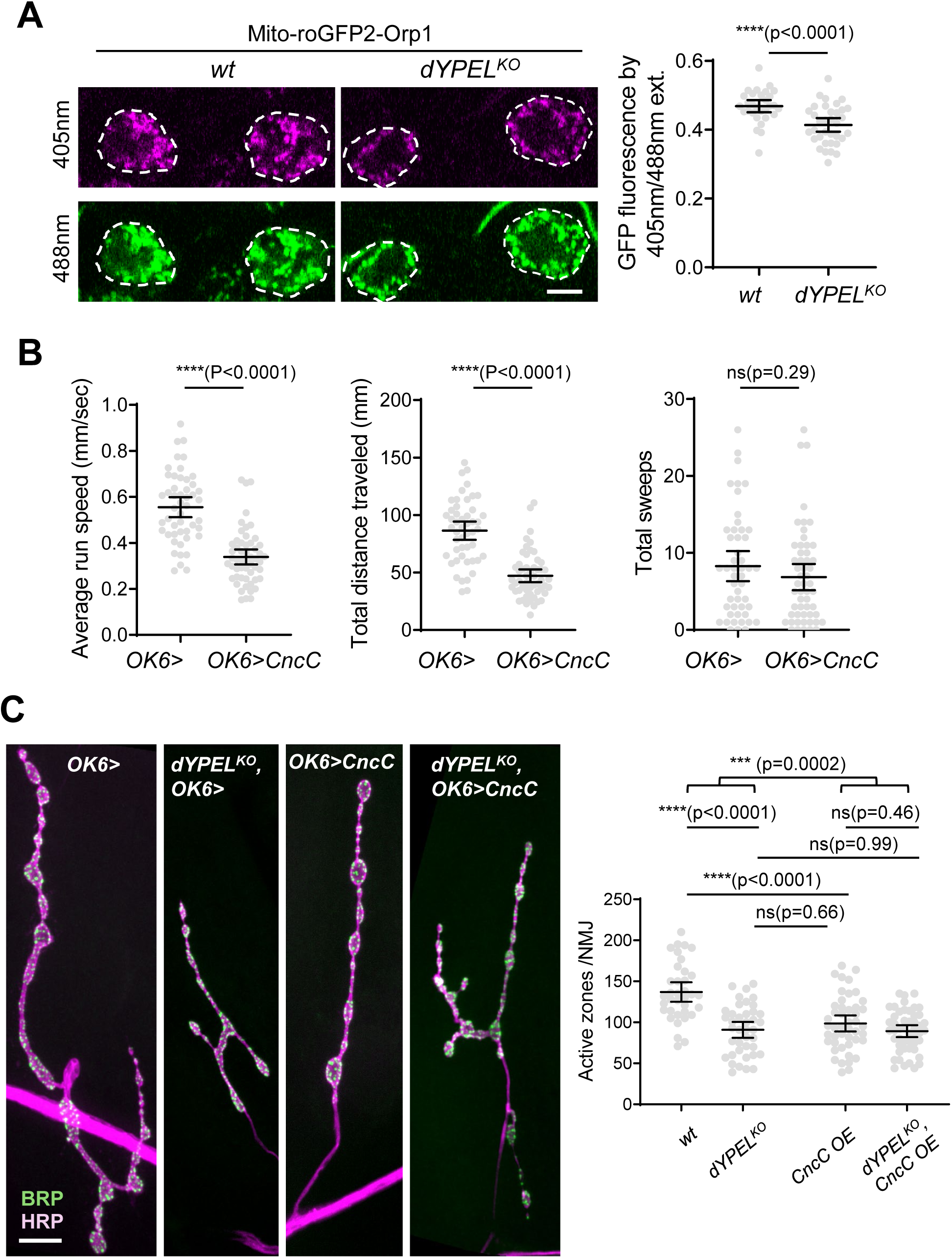
dYPEL regulates cellular antioxidant pathways. **(A)** *dYPEL* mutations caused a reduction in mitochondrial H_2_O_2_. Mitochondrial targeting sensor Mito-roGFP2-Orp1 was expressed in wt (*w^1118^*) and *dYPEL^KO^* larval motor neurons using *OK6-Gal4*. Representative images of roGFP green fluorescence from motor neuron cell bodies upon 405 nm (magenta) and 488 nm (green) excitation were shown. Samples were obtained from third-instar male larvae. Scale bar = 10 μm. The ratio of the green fluorescence obtained from 405 nm and 488 nm excitation was calculated and expressed as mean ± 95% CI. The genotypes and sample numbers were :*OK6>mito-roGFP2-Orp1* (*wt*; n = 30); *dYPEL^KO^* and *OK6>mito-roGFP2-Orp1* (n = 35). The two tailed Mann-Whitney test were used (U=235, p<0.0001). **(B)** Overexpression of CncC impairs locomotor function in *Drosophila* larvae. CncC was expressed in the larval motor neurons using *OK6-Gal4*. The driver line of *OK6-Gal4* only was used as a control. The third instar larvae were placed in a crawling behavior arena. Individual larva crawling was traced and quantified using a custom code to obtain the average run speed, the total distance traveled, and total number of sweeps. Data was presented as mean ± 95% CI. The genotypes and sample numbers were: the wild type control (*OK6>,* n = 49) and *OK6>*CncC (n = 54). The two tailed Mann-Whitney test were used (U=336, p<0.0001 for the average run speed; U=337, p<0.0001 for the total distance travelled; U=1163, p=0.2901 for the total sweeps). **(C)** CncC overexpression is epistatic with *dYPEL* mutations in neuromuscular synapse. CncC was expressed in wild type control and *dYPEL^KO^* larval motor neurons using *OK6-Gal4*. An anti-BRP antibody was used to visualize presynaptic active zones (green). HRP staining was used to visualize the type Ib boutons (magenta). Representative NMJ images of wild type control (*w^1118^, OK6>*), *dYPEL^KO^* (*dYPEL^KO^*, *OK6>*), CncC overexpression (*OK6>CncC*), and CncC overexpression in *dYPEL^KO^* (*dYPEL^KO^, OK6>CncC*). Scale bar = 10 μm. Samples were obtained from third-instar male larvae. The total number of BRP-positive puncta, or “Active zone” was quantitated from the type Ib NMJ images and expressed as mean ± 95% CI. The genotypes and sample numbers were: *OK6>* (n = 36); *dYPEL^KO^,OK6>* (n = 40); *OK6>CncC* (n = 46); *dYPEL^KO^*, *OK6>CncC* (n = 49). Two-way ANOVA (F(1, 167) = 15.01) followed by post hoc Turkey’s correction.

The C isoform of *Cap ‘n’ collar* (*CncC*) is the *Drosophila* ortholog of *Nrf2*. To test whether overactive Nrf2 signaling affects NMJ synaptic development and locomotion, we overexpressed CncC in motor neurons using *OK6-Gal4* and measured larva crawling behavior. The results showed a dramatic 39% reduction in the average running speed and a 45% decrease in the travel distance by CncC overexpression as compared to wild-type control (**Figure 5B**). No significant difference was observed in the head sweeps between control and CncC overexpression groups (**Figure 5B**). *OK6-Gal4* without CncC was used as a wild-type control. These behavioral phenotypes are highly reminiscent of those from *dYPEL* mutants **(Figure 1)**.

Next, we examined the effects of CncC overexpression on NMJ synaptic development. CncC was expressed in motor neurons under *OK6-Gal4* driver, and the NMJ was analyzed using anti-HRP and BRP antibodies. The results showed that CncC overexpression caused a significant 29% reduction in synaptic active zones compared to a control (**Figure 5C**), suggesting that excessive Nrf2/CncC activity impairs NMJ synapse development. Importantly, no additional reduction in active zones was observed when CncC was overexpressed in *dYPEL^KO^* motor neurons (**Figure 5C**), which supports a model wherein dYPEL and CncC function in a shared signaling pathway.

### dYPEL regulates synapse development through Nrf2/CncC pathway

A reduction in ROS levels in *dYPEL* mutants suggests an overactive Nrf2/CncC pathway. Furthermore, CncC overexpression phenocopied the reduction in synaptic active zones and the locomotion deficits seen in *dYPEL* mutants. These suggest that unregulated Nrf2/CncC activity is the root cause of *dYPEL* mutant phenotypes. Thus, we attempted to rescue *dYPEL* mutant phenotypes by reducing Nrf2/CncC activity. To this end, we used a knock-down approach. Cnc RNAi was expressed in wild-type and *dYPEL^KO^* larval motor neurons using *OK6-Gal4*. A transgene encoding RNAi for firefly luciferase (LUC-RNAi) was used as a negative control. The active zone in the NMJ and larval crawling behavior were measured.

Interestingly, we observed a significant increase in active zones in the NMJs from two *Cnc*-RNAi lines - a 38% and a 28% increase, while the third *Cnc*-RNAi line did not show a difference from the wild-type control. Although we do not know the implication of these increases, knocking down *Cnc* in *dYPEL^KO^* completely restored the active zones to their corresponding controls — *Cnc*-knockdown in wild-types (**Figure 6A and B**).

**Figure 6.**
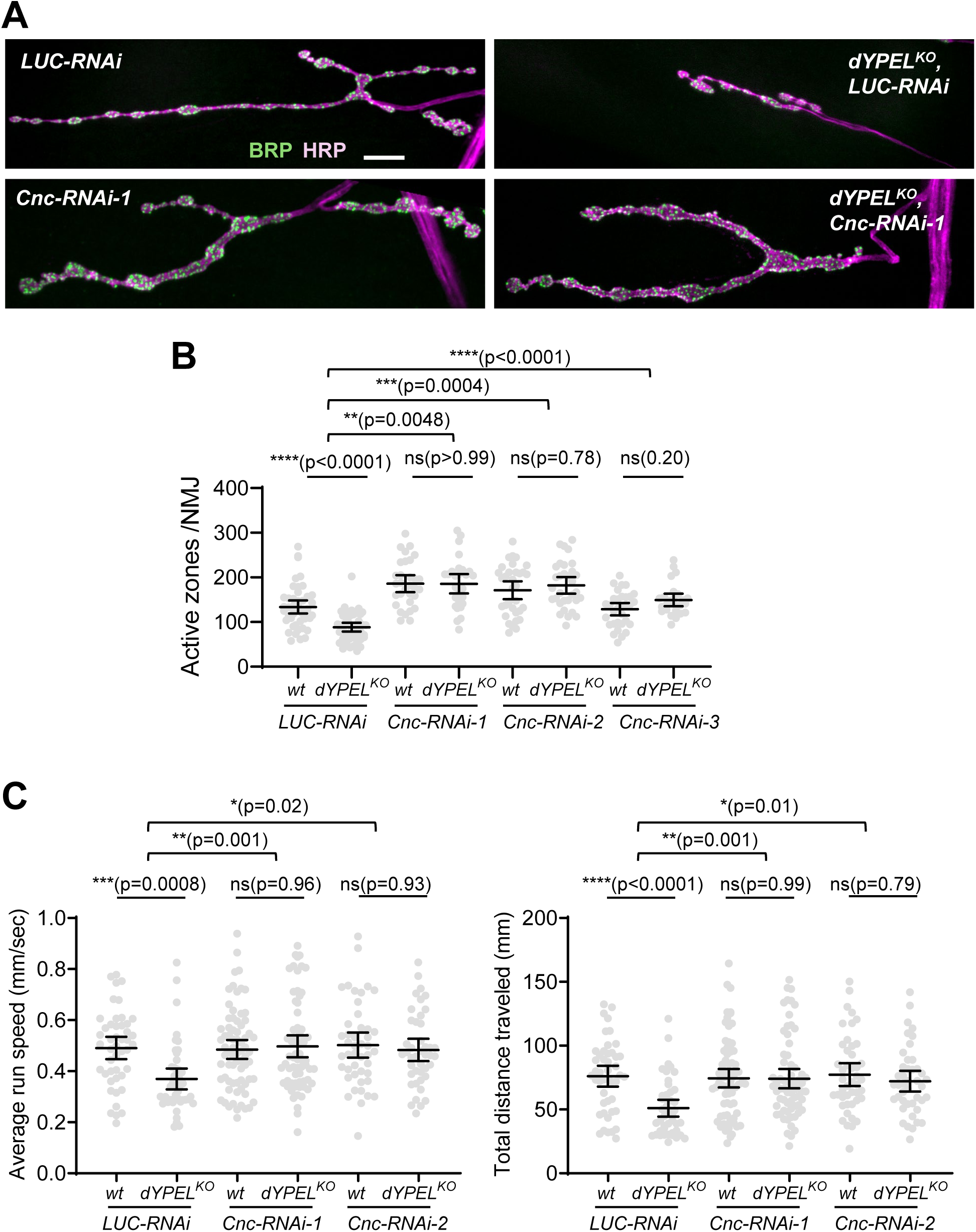
Cnc inhibition fully rescues the synaptic and locomotor deficits in *dYPEL mutants*. **(A) and (B)** *Cnc* knockdown rescues synaptic defects in *dYPEL* mutant NMJ. (A) A *Cnc-RNAi* (Cnc-RNAi-1) was expressed in wild type control and *dYPEL^KO^* larval motor neurons using *OK6-Gal4*. A RNAi for firefly luciferase (*LUC-RNAi*) was used as a negative control. An anti-BRP antibody was used to visualize presynaptic active zones (green). HRP staining was used to visualize the type Ib boutons (magenta). Representative NMJ images of wild type control (*LUC-RNAi*), *dYPEL^KO^* (*dYPEL^KO^*, *LUC-RNAi*), Cnc knockdown (*Cnc-RNAi*), and *Cnc* knockdown in *dYPEL^KO^* (*dYPEL^KO^, Cnc-RNAi*). Scale bar = 10 μm. Samples were obtained from third-instar male larvae. (B) Three different *Cnc-RNAi* (Cnc-RNAi-1, Cnc-RNAi-2, Cnc-RNAi-3) were expressed in wild type control and *dYPEL^KO^* larval motor neurons using *OK6-Gal4*. A RNAi for firefly luciferase (*LUC-RNAi*) was used as a negative control. The total number of BRP-positive puncta, or “Active zone” was quantitated from the type Ib NMJ images and expressed as mean ± 95% CI. The genotypes and sample number were: *OK6-Gal4*, UAS-Dcr-2, UAS-pLUZ-RNAi (n = 45); *dYPEL^KO^*, *OK6-Gal4*, UAS-Dcr-2, UAS-pLUZ-RNAi (n = 45); *OK6-Gal4*, UAS-Dcr-2, UAS-Cnc-RNAi line 1 (n = 31); *dYPEL^KO^*,*OK6-Gal4*, UAS-Dcr-2, UAS-Cnc-RNAi line 1 (n = 28); *OK6-Gal4*, UAS-Dcr-2, UAS-Cnc-RNAi line 2 (n = 33); *dYPEL^KO^*,*OK6-Gal4*, UAS-Dcr-2, UAS-Cnc-RNAi line 2 (n = 32); *OK6-Gal4*,UAS-Dcr-2/UAS-Cnc-RNAi line 3 (n = 32); *dYPEL^KO^*,*OK6-Gal4*, UAS-Dcr-2/UAS-Cnc-RNAi line 3 (n = 26). Two-way ANOVA was used to evaluate interactions between the LUC-RNAi groups and Cnc-RNAi groups pairwise followed by post hoc Turkey’s correction (F(1, 145) =8.209 for the control and Cnc-RNAi-1; F(1, 152) = 13.22 for the control and Cnc-RNAi-2; F(1,144) = 23.96 for the control and Cnc-RNAi-3). (C) *Cnc* knockdown rescues locomotor defects in *dYPEL* mutant. Two different *Cnc-RNAi* (Cnc-RNAi-1, Cnc-RNAi-2) were expressed in wild type control and *dYPEL^KO^* larval motor neurons using *OK6-Gal4*. A RNAi for firefly luciferase (*LUC-RNAi*) was used as a negative control. The third instar male larvae were placed in a crawling behavior arena. Individual larva crawling was traced and quantified using a custom code to obtain the average run speed and the total distance traveled. Data was presented as mean ± 95% CI. Genotypes and sample numbers were: *OK6-Gal4*, UAS-Dcr-2, UAS-pLUZ-RNAi (n = 46); *dYPEL^KO^*,*OK6-Gal4*, UAS-Dcr-2, UAS-pLUZ-RNAi (n = 45); *OK6-Gal4,* UAS-Dcr-2, Cnc-RNAi-1, (n = 72); *OK6-Gal4*, UAS-Dcr-2, UAS-Cnc-RNAi-2 (n = 45); *dYPEL^KO^*,*OK6-Gal4*, UAS-Dcr-2, Cnc-RNAi-1 (n = 67); *dYPEL^KO^*, *OK6-Gal4*, UAS-Dcr-2, UAS-Cnc-RNAi-2 (n = 43). Two-way ANOVA was used to evaluate interactions between the LUC-RNAi groups and Cnc-RNAi groups pairwise followed by post hoc Turkey’s correction. The average run speed (F(1, 226) = 9.825 for the control and Cnc-RNAi-1; F(1, 173) = 5.363 for the control and Cnc-RNAi-2). The total distance traveled (F(1, 226) = 10 for the control and Cnc-RNAi-1; F(1, 175) = 6.252 for the control and Cnc-RNAi-2).

For larva crawling behavior, two *Cnc*-RNAi fly lines were used. Similarly, LUC-RNAi was used as a negative control. *Cnc*-RNAi was expressed in motor neurons using *OK6-Gal4* driver. Consistent with the rescue of the active zones in the NMJ, knocking down *Cnc* completely rescued the locomotor defects in *dYPEL^KO^* larvae (**Figure 6C**). While dYPEL^KO^ significantly impaired larva locomotion, the average run speed and the travel distance of *dYPEL^KO^* larvae expressing *Cnc*-RNAi were indistinguishable from those of wild-type control, which was a 34%/31% increase in the average run speed and a 45% and a 42% increase in the travel distance by the two *Cnc*-RNAi lines respectively. Both the average run speed and the total distance traveled in two *Cnc*-RNAi expressions were not different from those in the control group (**Figure 6C**).

Taken together, these findings strongly suggest that YPEL regulates NMJ development and locomotor activity by modulating Nrf2/CncC activity.

In summary, our study proposes a model where YPEL suppresses p62/Ref(2)P bodies, ensuring proper suppression of Nrf2/CncC. However, when YPEL function is compromised, p62/Ref(2)P bodies are increased, leading to unregulated Nrf2/CncC activity. Overactive Nrf2/CncC suppresses ROS levels below the physiological threshold, resulting in defective NMJ synaptic development and motor functions (**Figure 7**).

**Figure 7.**
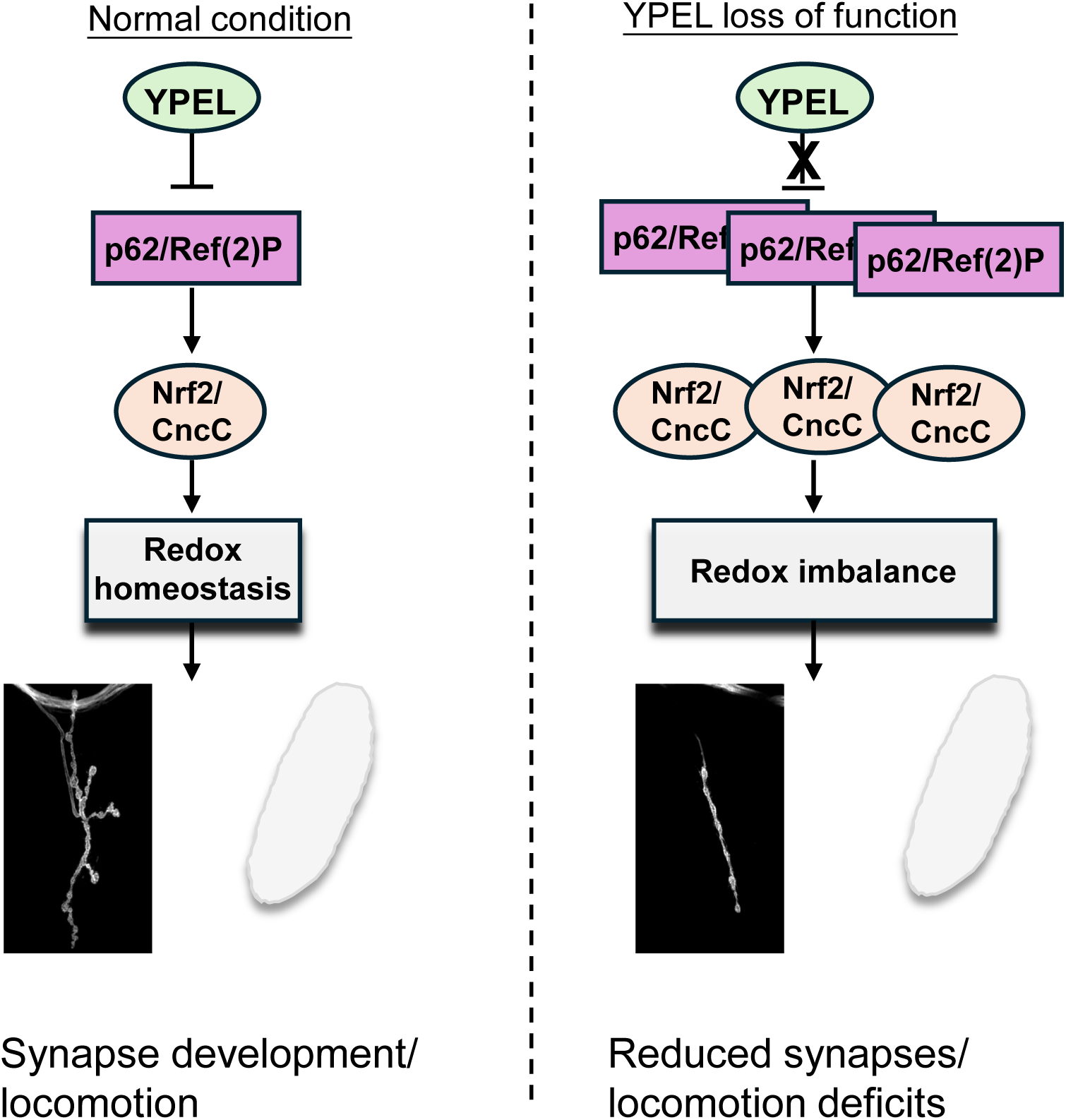
The model of YPEL function on synapse development and locomotor activity. YPEL inhibits p62/Ref(2)P body formation, thus suppresses Nrf2/CncC activity under normal conditions, which ensures proper amount of reactive oxygen species for synapse development. However, under *YPEL* loss-of-function conditions, p62/Ref(2)P bodies increase, leading to heightened Nrf2/CncC activity, which scavenges reactive oxygen species below physiological thresholds. This leads to a reduction in neuromuscular synapses and locomotion deficits.

## Discussion

*YPEL* genes are present in all eukaryotes and highly conserved, which suggests that YPEL proteins perform fundamental cellular functions. Furthermore, a gene mutation in *YPEL3* — one of the *dYPEL* homologs in humans, was found in a patient with locomotor deficits, emphasizing a potential role of YPEL3 in human motor functions. However, the molecular and cellular functions of YPEL proteins are poorly understood. Here, we demonstrate that YPEL negatively controls neuronal p62 bodies to suppress Nrf2 antioxidant pathway. We showcase that this regulation is required for neuromuscular synapse development and locomotion. Importantly, we demonstrate that suppressing Nrf2 activity is sufficient to fully rescue the deficits in both neuromuscular synapses and motor activity in *YPEL* mutants. Together, we demonstrate a previously unknown cellular function of YPEL in synapse development via an antioxidant pathway.

### The role of YPEL in locomotion and NMJ/synapse development

Various deletions and duplications in the chromosome region that contains *YPEL3* are associated with multiple brain disorders [13–19]. A human study suggests that YPEL3, one of the *Drosophila YPEL* homologs in humans, is necessary for proper motor activity [20]. In zebrafish, a morpholino-mediated targeting of *YPEL3* altered brain structures [19]. Another study characterized defects in glial cell development in *YPEL3* mutant Zebrafish, which coincided with altered swimming pattern [20]. In the present study, we showed that both the frameshift and genetic null *dYPEL* mutations resulted in crawling behavior deficits as manifested as slower and shorter movement bouts (**Figure 1**). We further showed that *dYPEL* mutant larvae display smaller NMJ sizes and reduced synapses (**Figure 1**). Although it is possible that the observed locomotor deficits may be secondary to defective nervous system development, we suggest the reduction in active zones is the main cause of defective locomotor activity. First, larva crawling was measured in the absence of any added sensory stimulation under complete darkness. Secondly, *dYPEL* mutants did not show any changes in sweeping behavior. Sweeping behavior is associated with the internal state of an animal and thus can be used as an indirect measurement of the nervous system functioning. Moreover, the locomotor deficits could be completely rescued by motor neuron-specific knock-down of *Cnc*, which also rescued NMJ synapses in *dYPEL* mutants (**Figure 6**). Taken together, these suggest that the locomotor defects are highly associated with NMJ function under our experimental conditions. Although we did not measure the NMJ transmission efficacy, reduced NMJ sizes and reduced active zones strongly suggest the observed locomotor deficits stem from reduced synaptic transmission in the NMJ. This is consistent with our previous work, which showed reduced synaptic transmissions in *dYPEL* mutant *Drosophila* larval sensory circuits [21].

### YPEL regulation on p62 bodies for NMJ/synapse development

How does YPEL control synapse development? A study has shown that YPEL3 positively regulates Glycogen synthase kinase-3 (GSK3) in a tumor cell spreading model [40]. GSK3 has well-known functions in synaptic plasticity [41], which suggests that YPEL - GSK3 pathway may be responsible for NMJ synapse development. Mutations in fly GSK3 resulted in synaptic overgrowth in larva NMJ [42]. On the other hand, our study demonstrated that *dYPEL* loss caused smaller NMJ. Thus, we believe that YPEL – GSK3 pathway is unlikely responsible for neuromuscular synapse development.

A proteome-wide screening identified Ref(2)P as one of the potential dYPEL binding proteins in *Drosophila* although functional relationships were not known [31]. Abnormal accumulation of p62-containing cytoplasmic bodies was observed from multiple human disorders including Alzheimer’s and Parkinson’s disease [22]. Moreover, a number of *p62* missense mutations are associated with multiple diseases including amyotrophic lateral sclerosis and frontotemporal lobar degeneration [23]. These suggest a potential involvement of p62 or p62 bodies in the pathogenesis of these disorders. We found that dYPEL proteins form physical complex with Ref(2)P and dramatically decrease Ref(2)P bodies when overexpressed (**Figure 2**). Consistently, *dYPEL* mutations caused an increase in neuronal p62 bodies. Importantly, normalizing p62 bodies in *dYPEL* mutant significantly alleviated both synaptic and locomotor deficits (**Figure 3**). These strongly suggest that YPEL negatively regulates p62 bodies, and that the increased p62 bodies are causative for *dYPEL* mutant phenotypes in synapse development and locomotion. To our knowledge, our study provides the first causal evidence of neuronal defects by increased p62 bodies. A similar mechanism might be in place for disease pathogenesis by abnormal accumulation of p62 bodies seen in neurodegenerative disorders.

p62 oligomers generate cytoplasmic proteinaceous bodies along with polyubiquitinated protein aggregates for autophagic degradation [26]. Increasing autophagic activity by Atg1 overexpression was sufficient for reducing Ref(2)P bodies in *dYPEL* mutants. However, interestingly, it did not restore defective synapse development (**Figure 4**). Rather, Atg1 overexpression caused dramatic synaptic defects. One potential explanation is that our experimental settings may have reduced p62 bodies below the physiological threshold and induced p62 loss of function. Consistent with this idea, *p62*-loss was found in individuals with early-onset neurodegeneration symptoms that are characterized by ataxia and cognitive impairment [24, 25]. Moreover, *p62* knock-out mice developed Alzheimer-like phenotypes with anxiety, depression, and loss of working memory [43]. Together, these suggest that keeping proper amounts of p62 bodies is essential for proper synapse development. Whether the loss and the gain of p62 bodies trigger the same molecular pathway is not known and requires important future investigations. Another interesting point is that autophagy is involved in various aspects of neuronal functions [28] and in fly NMJ development [44]. Thus, whether changes in autophagic activity affect synapses and NMJ development through p62 bodies under physiological and pathological conditions will be an interesting future research question.

*YPEL* genes encode relatively small protein products of about 12 - 15kDa. And their molecular functions are highly elusive. A recent study demonstrated that YPEL5 – an ortholog of *Drosophila Yipee*, inhibits an E3-ubiquitin ligase complex, WDR26-CTLH, by acting as a substrate mimicry [45]. Their crystal structure analysis reveals that both the N-terminal and the rigid structured region engages electrostatic and molecular complementarity interaction for YPEL5-WDR26 binding [45]. While the amino acid residues that are involved in molecular interaction between YPEL5 and WDR26 are 100% conserved between human YPEL5 and Yippee, those amino acid sequences display limited homology in other YPEL homologs including dYPEL. Whether YPEL employs the same or a similar molecular function as YPEL5, or functions through distinct molecular mechanisms for regulating p62 bodies warrant further investigations.

### The role of Nrf2 and cellular redox in YPEL-mediated NMJ/synapse development

We found reduced levels of ROS in *dYPEL* mutant motor neurons (**Figure 5**). p62 bodies participate in diverse cell signaling events through modular motifs and domains [27]. One of these signaling events triggers cellular antioxidant response via Nrf2 activation [27]. Overactivation of *Drosophila* Nrf2, CncC phenocopied the synaptic and locomotor deficits seen in *dYPEL* mutants. Importantly, knocking down *Cnc* in *dYPEL* mutant motor neurons fully rescued the NMJ and locomotor defects (**Figure 6**).

How might hyperactive Nrf2/CncC or antioxidant response negatively affect synapse development? Abnormal levels of oxidative stress are a hallmark of many neurodegenerative disorders. Consistent with this idea, reduced Nrf2 activation is associated with Alzheimer and Parkinson diseases [46]. Nrf2 promotes neuronal differentiation of a neuroblastoma cell line [47]. Nrf2 overexpression promoted neuronal survival in neurodegenerative conditions in mouse photoreceptors and retinal ganglion cells [48]. Interestingly, Nrf2 knock-out mice exacerbate cognitive deficits in Alzheimer’s disease model [49, 50] and in cognitive dysfunction model [51], suggesting a potential role of Nrf2-dependent antioxidant response in protecting neuronal synapses.

On the other hand, neuronal function also relies on certain ROS levels. For example, ROS levels are increased by neural activity, which in turn promotes synaptic bouton formation in fly NMJ [52] and inhibits postsynaptic glutamate receptor mobilization for synaptic homeostasis [53]. Interestingly, epigenetic repression of *Nrf2* was reported in mouse cortical neurons in vitro and in vivo [38, 54], which was necessary for neurite and synapse formation [38]. A recent report tested the role of presynaptic mitochondrial ROS more directly and has shown an increased synaptic active zones [55], which might explain the reduced synaptic active zones by CncC overexpression and *dYPEL* mutations. It will be an important future investigation to identify direct ROS targets in synapse regulation.

### Potential role of YPEL regulation on p62/Nrf2 pathway beyond synapse development

A number of research points to the potential role of *YPEL* genes in cellular senescence and tumor suppression [5–8]. *YPEL3* is a p53-regulated gene that induces cellular senescence and apoptosis [6]. Downregulation of YPEL3 has been observed in various human tumors, such as colon, lung, and ovarian cancers [56]. Moreover, knocking down *YPEL* genes in clear cell renal cell carcinoma cell lines led to increased cell proliferation, migration, and invasion [57]. Interestingly, elevated levels of p62 have been observed in various cancers and are often associated with poor prognosis. For example, high p62 expression correlates with increased metastasis in colon cancer [58]. In pancreatic cancer, elevated p62 levels are associated with reduced survival rates post-tumor resection [59]. Increased p62 expression is linked to distant metastasis and unfavorable prognosis in breast cancer [60]. In light of our findings that YPEL negatively regulates p62 bodies (**Figure 2**), these raise an interesting possibility that YPEL acts as a tumor suppressor by inhibiting p62 bodies. Nrf2 is also implicated in tumorigenesis but with a highly context-dependent manner [61]. Whether p62 suppression is the mechanistic basis for the tumor suppressor role of YPEL remains to be determined and will be an interesting future investigation.

In summary, our study revealed the novel findings of YPEL on p62 regulation and provides a potential involvement of heightened Nrf2 antioxidant response in *YPEL* loss-induced NMJ/synapse pathogenesis and locomotor deficits. Due to the potential roles of Nrf2 in neurodegeneration and cancer, various attempts are in progress to develop Nrf2 modulators in clinical settings [62, 63]. We envision that our findings will inform an essential rationale for using Nrf2 modulators for YPEL-related disorders.

## MATERIALS AND METHODS

### Drosophila melanogaster strains

*Drosophila* strains were kept under standard condition at 25 °C in a humidified chamber. The following strains were used in this study: *w^1118^*(3605), *dYPEL^T1-8^* and *dYPEL^KO^* [21]*, Ref(2)P^OD2^* (99019), *Ref(2)P^OD3^*(99020), *UAS-Atg1* (51654), *OK6-Gal4* (64199), *UAS-cncC* (95246), *UAS-cnc-RNAi* (25984), *UAS-cnc-RNAi* (32863), *UAS-cnc-RNAi* (40854), *UAS-pLUC-RNAi* (31603), *UAS-Dcr-2* (24645), *ppk-Gal4* [34], *UAS-mito-roGFP2-Orp1* (67667).The numbers in parentheses indicate the stock numbers from the Bloomington *Drosophila* Stock Center.

### DNA constructs

The UAS-Ref(2)P-GFP construct was generated using the In-Fusion Cloning Kit (Takara Bio USA, San Jose, CA). The Ref(2)P cDNA open reading frame (isoform B) was purchased from GenScript Biotech (Piscataway, NJ). *GFP* cDNA was recovered from a UAS-GFP construct [64]. The coding sequences from *Ref(2)P* and *GFP* were amplified by PCR using CloneAmp HiFi PCR Premix (Takara Bio USA, San Jose, CA). The primers used were TGAATAGGGAATTG GGAATTATGCCGGAGAAGCTGTTGAAA and GTTGCGGTTCTGCGATACGT for Ref(2)P, and ACGTATCGCAGAACCGCAACGGAGGAGGAATGGTGAGCAAGGGCGAGG and GGTACCCTCGAGCCGCGGCCTTACTTGTA CAGCTCGTCCATGCC for GFP. The pUASTattB plasmid was linearized using NotI and EcoRI restriction sites.

To generate a C-terminally Myc-tagged dYPEL, the coding sequence of dYPEL was PCR amplified and inserted into pUASTattB plasmid using NotI and XbaI restriction sites. The Myc sequences were added as the flanking sequences in the reverse primer. The primers used were the forward TAAGCGGCCGCATGGTGAAGACGTTTCAGGCCTATCTACCGTCC and the reverse AGTCTAGATTACAGATCCTCTTCAGAGATGAGT TT CTGCTCTCCTCCTGCGTCC CAGCCGTTCTCCTTGATCATGTGC.

### Cell culture, transfection, co-immunoprecipitation and Western blot analysis

*Drosophila* Schneider 2 (S2) cells were cultured in Schneider’s Medium (Thermo Fisher Scientific, Waltham, MA) supplemented with 10% fetal bovine serum (Sigma-Aldrich, Burlington, MA) at 27 °C in a humidified incubator. Cells were transfected with the expression plasmids along with *tubulinP*-Gal4 using polyethylenimine (Polysciences, Warrington, PA) as previously described [65].

Forty-eight hours after transfection, S2 cells were harvested by centrifugation. Cells were lysed in Lysis buffer (20 mM Tris/pH7.2, 150 mM NaCl,1% NP-40, 0.1 mM DTT) containing the Halt^TM^ protease inhibitor cocktail EDTA-free (Thermo Fisher Scientific, Waltham, MA), 1 mM Sodium vanadate (Thermo Fisher Scientific, Waltham, MA), and 5 mM sodium fluoride (Alfa Aesar, Wall Hill, MA) on ice for 30 min. The nuclear fraction was removed by centrifugation at 1000 x g for 5 min. The post nuclear fraction was further centrifuged at 20,000 x g for 30 min at 4 °C. The resulting supernatant was incubated with the GFP Selector (NanoTag Biotechnologies, Göttingen, Germany) for 1 hour at 4 °C on a head-over-tail rotator. The protein bound GFP selector was recovered by centrifugation for 10 sec at 1000 x g at 4 °C. The supernatant was discarded, and the GFP Selector was resuspended in 1 ml Lysis buffer and washed twice in Lysis buffer. Then, 20 μl of SDS sample buffer was directly added to the GFP Selector followed by heating at 95 °C for 5 min, then centrifuged at 20,000 x g for 5 min immediately. The resulting eluents were used for subsequent Western blot analysis.

For preparing the larval brain lysates, the central nervous system including the ventral nerve cords was isolated from the wandering 3^rd^ instar larvae in ice-cold phosphate-buffered saline (PBS). Larval brains were immediately lysed in 2XSDS-PAGE sample buffer by vortexing for 30 sec, followed by heating at 95 °C for 5 min, and then centrifuged at 20,000 x g for 5 min.

For preparing the adult heads, 2–5 day old adult fly heads collected in ice-cold PBS and stored at −20 °C until use. 2XSDS-PAGE sample buffer was added to fly heads, followed by hand pestle homogenization, then heating at 95 °C for 5 min, and then centrifuged at 20,000 x g for 5 min.

Western blot analysis was performed as previously described [65]. Primary antibodies used were Chicken polyclonal anti-GFP (Aves Labs, Tigard, OR), Chicken polyclonal anti-Myc (Aves Labs, Tigard, OR), Rabbit polyclonal anti-Ref(2)P (Abcam, Boston, MA), Rat monoclonal anti-Elav (DSHB, Iowa city, IA), Rabbit polyclonal anti-ATG8 (Sigma-Aldrich, Saint Louis, MO). Secondary antibodies used were from Jackson ImmunoResearch (West Grove, PA): Donkey HRP conjugated anti-chicken IgG, Donkey HRP conjugated anti-rabbit IgG, and Donkey HRP conjugated anti-rat IgG.

### Immunostaining and imaging

The wandering 3^rd^-instar larvae were dissected in PBS and were fixed with fixation solution for 45 min at room temperature. Samples were washed three times with PBST (PBS containing 0.1% Triton X-100). The samples were then blocked in blocking solution (PBST with 3% normal Donkey serum) for 1 hour at room temperature. Samples were incubated with primary antibodies overnight at 4 °C. Samples were washed three times with PBST before being incubated with fluorophore-conjugated secondary antibodies overnight at 4 °C. All antibody solutions were prepared in blocking solutions. The samples were mounted using Prolong Gold Antifade reagent (Thermo Fisher Scientific, Waltham, MA).

Twenty-four hours after transfection, S2 cells were harvested by gentle pipetting and plated on concanavalin-A (ConA) coated coverslips. A ConA-coated coverslip was prepared using a 0.5 mg/ml ConA solution (MP Biomedicals, Solon, OH) in autoclaved Milli-Q water. The solution was filtered through a 0.22 μm syringe filter before use. The coverslip was placed on a piece of Parafilm, and 200 μL of the ConA solution was evenly spread across its surface, followed by incubation for 30 minutes at room temperature. Excess solution was removed by carefully lifting the coverslip. The coverslip was then air-dried in a culture hood for 15 minutes and subsequently sterilized under UV light for 15 minutes before use. Cells were allowed to spread for an additional 24 hours in the incubator. Subsequently, cells were rinsed with PBS and fixed with fixation solution (4% formaldehyde in PBS) for 45 minutes at room temperature. The coverslips were washed three times with PBST, blocked for 30 min, and then incubated with primary antibodies for 1 hour at room temperature. The coverslips were washed three times with PBST before being incubated with secondary antibodies for 1 hour at room temperature. Finally, the coverslips were washed three times with PBST before being mounted with Prolong Gold Antifade reagent (Thermo Fisher Scientific, Waltham, MA).

Antibodies used are Chicken polyclonal anti-myc (Aves Labs, Tigard, OR), Rabbit polyclonal anti-Ref(2)P (Abcam, Boston, MA), and Mouse monoclonal anti-BRP (DSHB, lowa city, IA). Secondary antibodies were from Jackson ImmunoResearch (West Grove, PA): Cy2 or Cy5-conjugated Goat anti-HRP, Cy2-conjugated AffiniPure Donkey anti-chicken IgY, and Cy3-conjugated AffiniPure Donkey anti-rabbit IgG.

Fluorescence microscope imaging was performed using a custom-built spinning disk confocal microscope equipped with a 63X immersion oil lens with a 0.3 μm-step size. Images were acquired with a 16-bit image depth. For neuromuscular junction (NMJ) analysis, Type Ib NMJ on muscle 4 from the abdominal segments 2 to 5 were imaged. For Ref(2)P analysis, C4da neurons from the abdominal segments 4 to 6 were imaged. Resulting 3D image stacks were projected into 2D images using a maximum projection method. For the Ref(2)P puncta analysis of C4da neurons, a single image slice containing the maximum number of Ref(2)P puncta was selected for analysis. For a given analysis, essentially the same imaging settings were used.

### Measurement of H_2_O_2_ in vivo

To measure hydrogen peroxide levels in vivo, third-instar larvae were removed from the food vial, briefly rinsed with distilled water before being transferred to PBS containing 2 mM N-ethylmaleimide (NEM)(Thermo scientific, Rockford, IL) and immediately dissected. NEM alkylates free sulfhydryl groups on roGFP thus, preventing further oxidation during sample preparation and imaging [39].

To achieve full alkylation of cysteine sulfhydryl groups, dissected larvae were further incubated in 2 mM NEM solution for 10 minutes at room temperature (RT). Samples were briefly rinsed with PBS to remove excess NEM before being fixed in the fixation solution for 15 minutes at room temperature. Then, the samples were washed twice with PBS and mounted using Prolong Gold Antifade reagent. The samples were kept in the dark for 24 hours at 4 °C before being imaged.

The roGFP imaging was performed using a Leica SP8 laser scanning confocal microscope equipped with a 63X immersion oil lens. Green fluorescence emission was sequentially acquired from 510 – 530 nm using 405 nm and then 488 nm laser excitation with a step size of 0.3 μm from the motor neuron cell bodies. The motor neuron cell bodies were identified from the ventral nerve code segments that correspond to A4-A6. For the mean fluorescence measurement, a single image slice displaying clear/maximum GFP signals were selected, followed by background fluorescent subtraction. The numerical values for ratiometric analysis were obtained by dividing the mean fluorescence intensities from 405 nm excitation by those obtained from 488 nm excitation.

### Neuromuscular junction, BRP and Ref(2)P puncta image analysis

The NMJ morphology was analyzed using the ImageJ plugin “*Drosophila NMJ Morphometrics*” developed by Castells-Nobau et al [66]. First, NMJ images were processed using the sub-macro “convert to stack” to create Z-projections and hyperstacks. The sub-macro “define ROI” was then applied to define the NMJ of interest from the anti-HRP staining image channel. Subsequently, the sub-macro “Analyze” was used to quantify NMJ features with the following parameters: rolling ball radius: 20; minimum bouton size: 150 pixel; small particles size: 150 pixel; find maxima noise tolerance 50; Scale-pixels:1; Scale-distance in micrometers: 0.0984; NMJ outline threshold: RenyiEntropy; Skeleton threshold: Li.

For active zone analysis, BRP-positive puncta were identified and quantified. First, “Filters>Gaussian Blur” function [67] in ImageJ was applied to minimize background noise. Each image was duplicated to create two versions: image 1 with sigma 1, and image 2 with sigma 2. The “Image Calculator” function [68] was then used to subtract image 2 from image 1. The size and circularity thresholds were set to 10 pixels and 0.2, respectively. BRP puncta were counted using the “Analyze Particles” function in ImageJ (National Institutes of Health, RRID:SCR_003070). The intensity threshold was adjusted for individual images to ensure that all visibly identified puncta were counted. The region of interest was defined based on the HRP channel.

Ref(2)P puncta analysis was done essentially identical as BRP puncta analysis except that the region of interest was defined by the C4da cell body labelled with anti-HRP antibody, and that the intensity threshold was manually adjusted for a given experiment (the control and experimental groups), not for individual images.

### Larval locomotion behavioral assay

The larval crawling analysis was done as previously described [30]. Third-instar larvae were collected from food using a high-density (15 %) sucrose solution, washed four times with distilled water before placing in the behavioral arena. Larval crawling behavior was recorded in the dark with a monochrome USB 3.0 CCD camera fitted with an IR long-pass 830 nm filter and 8 mm F1.4 C-mount lens. An infrared (850 nm) LED light was used for illumination. The video was recorded for 3 minutes. A total of 20 larvae were used per an assay, with ten assays performed for each group on different days. The control and experimental groups were paired in a behavioral session in an alternative order with three to four assays performed a day. All assays were combined for statistical analysis. In addition, to minimize the potential behavioral variation by circadian rhythms, all experiments were performed at approximately the same time of the day within each session. The behavioral assays were conducted in a room maintained at 25 °C and 45 % humidity.

The behavioral parameters were calculated from the individual larva tracing using custom routines written in MATLAB (MathWorks Inc., Natick, MA, RRID:SCR_001622) [30]. Runs were defined as continuous periods of forward movement. Run length and Run speed were determined through the analysis of runs. Sweeps were defined when the larva was at a stop and the angle between the head and tail of the larva was greater than 20 degrees. A head sweep occurs when larvae perform of their head part of behavior such as searching for food, exploration, social interaction, or defense, among others.

### Experimental design and statistical analysis

All statistical analyses were performed in GraphPad Prism software (version 7.04, RRID:SCR_002798). For comparisons between two groups, the Mann-Whitney test was employed. One-way ANOVA followed by post hoc Turkey’s multiple comparison test was used for experiments involving three groups. Two-way ANOVA with post hoc multiple comparison using Turkey’s correction was used for genetic interaction experiments. Data was presented as mean ± 95% CI except for the Western blot data, which was presented as mean ± SEM. The statistical significance was set to * *p* < 0.05, \*\**p* < 0.01, \*\*\**p* <0.001, and \*\*\*\**p* < 0.0001.

## Acknowledgments

Research reported in this study used the Cellular and Molecular Imaging Core facility at the University of Nevada Reno.

---

**Figure 3 supplementary -1.**
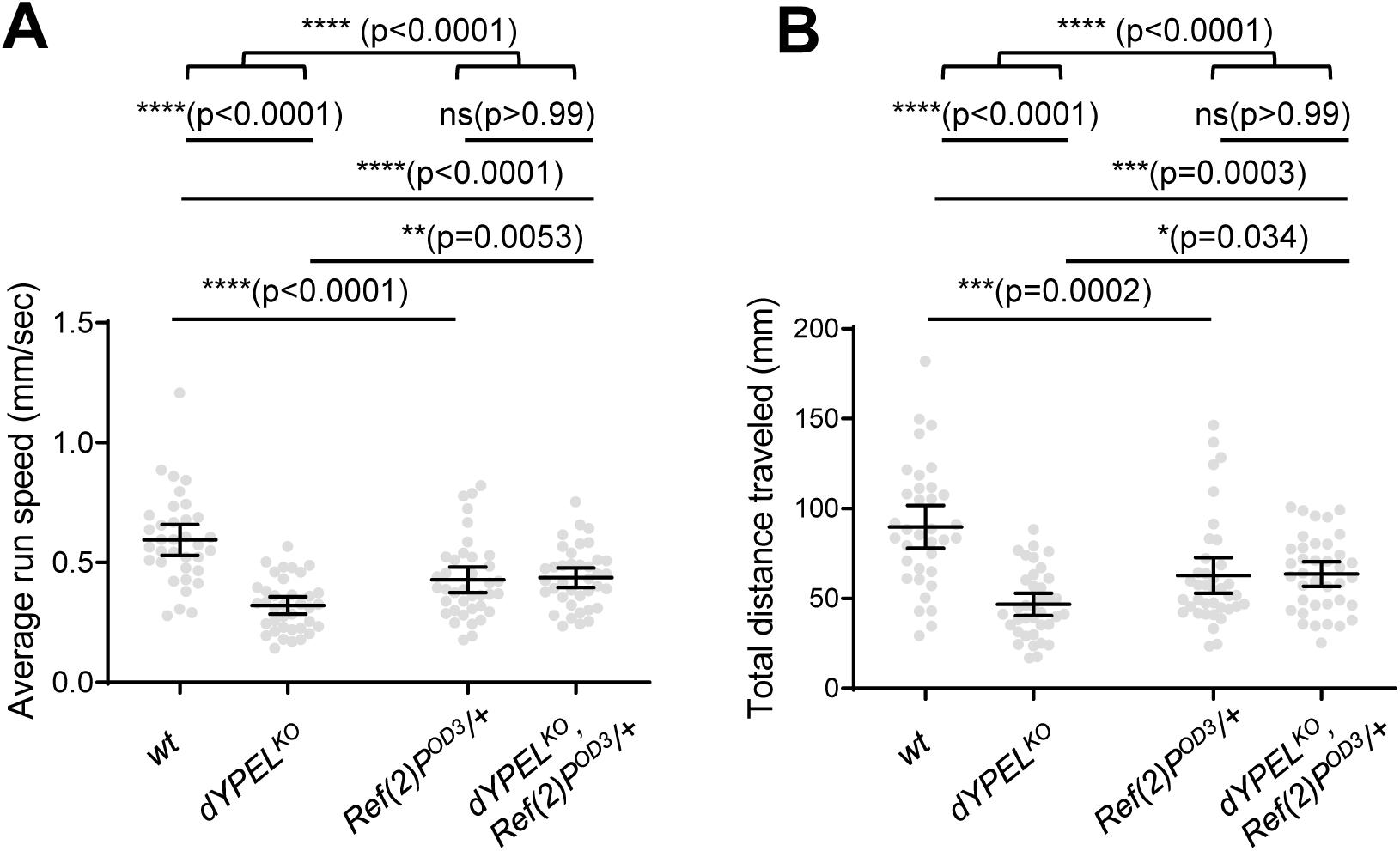
One copy of *Ref(2)P^OD3^* alleviated locomotor deficits in *dYPEL^KO^* The third instar male larvae were placed in a crawling behavior arena. Individual larva crawling was traced and quantified using a custom code to obtain the average run speed and the total distance traveled. Data was presented as mean ± 95% CI. The genotypes and sample numbers were *w^1118^* (n = 35); *dYPEL^KO^* (n = 37); *Ref(2)P^OD2^/+* (n = 38); *dYPEL^KO^*, Ref(2)P^OD2^/+ (n = 38). Average run speed: Two-way ANOVA (F(1, 144) = 33.25) followed by post hoc Turkey’s correction. Total distance travelled: Two-way ANOVA (F(1, 144) = 25.08) followed by post hoc Turkey’s correction.

## Reference

1. Hosono K, Sasaki T, Minoshima S, Shimizu N. Identification and characterization of a novel gene family YPEL in a wide spectrum of eukaryotic species. Gene. 2004;340(1):31–43. doi: 10.1016/j.gene.2004.06.014. PMID: 15556292

2. Han J, Shin J, Lee Y, Kim K. Distinct roles of the YPEL gene family in development and pathogenicity in the ascomycete fungus Magnaporthe oryzae. SCIENTIFIC REPORTS. 2018;8. doi: 10.1038/s41598-018-32633-6. PMID: 30262874

3. Tan TY, Gordon CT, Miller KA, Amor DJ, Farlie PG. YPEL1 overexpression in early avian craniofacial mesenchyme causes mandibular dysmorphogenesis by up-regulating apoptosis. Developmental Dynamics. 2015;244(8):1022–30. doi: 10.1002/dvdy.24299. PMID: 26061551

4. Farlie P, Reid C, Wilcox S, Peeters J, Reed G, Newgreen D. Ypel1: a novel nuclear protein that induces an epitheliallike morphology in fibroblasts. Genes to Cells. 2001;6(7):619–29. doi: 10.1046/j.1365-2443.2001.00445.x. PMID: 11473580

5. Xu J, Tang M, Lu Z, Song Y, Zhang K, He R, et al. A novel role for YPEL2 in mediating endothelial cellular senescence via the p53/p21 pathway. MECHANISMS OF AGEING AND DEVELOPMENT. 2023;211. doi: 10.1016/j.mad.2023.111803.PMID: 36963468

6. Kelley KD, Miller KR, Todd A, Kelley AR, Tuttle R, Berberich SJ. YPEL3, a p53-Regulated Gene that Induces Cellular Senescence. Cancer Research. 2010;70(9):3566–75. doi: 10.1158/0008-5472.can-09-3219. PMID: 20388804

7. Singh A, Thakur M, Singh SK, Sharma LK, Chandra K. Exploring the effect of nsSNPs in human YPEL3 gene in cellular senescence. Scientific Reports. 2020;10(1). doi: 10.1038/s41598-020-72333-8. PMID: 32943700

8. Tuttle R, Miller K, Maiorano J, Termuhlen P, Gao Y, Berberich S. Novel senescence associated gene, YPEL3, is repressed by estrogen in ER plus mammary tumor cells and required for tamoxifen-induced cellular senescence. INTERNATIONAL JOURNAL OF CANCER. 2012;130(10):2291–9. doi: 10.1002/ijc.26239. PMID: 21671470

9. Oki K, Plonczynski M, Gomez-Sanchez E, Gomez-Sanchez C. YPEL4 modulates HAC15 adrenal cell proliferation and is associated with tumor diameter. MOLECULAR AND CELLULAR ENDOCRINOLOGY. 2016;434(C):93–8. doi: 10.1016/j.mce.2016.06.022. PMID: 27333825

10. Mattebo A, Sen T, Jassinskaja M, Pimkova K, Gonzalez-Albo IP, Alattar AG, et al. Yippee like 4 (Ypel4) is essential for normal mouse red blood cell membrane integrity. Scientific Reports. 2021;11(1). doi: 10.1038/s41598-021-95291-1. PMID: 34354145

11. Deng Y, Han X, Chen H, Zhao C, Chen Y, Zhou J, et al. Ypel5 regulates liver development and function in zebrafish. JOURNAL OF MOLECULAR CELL BIOLOGY. 2023;15(3). doi: 10.1093/jmcb/mjad019. PMID: 36948605

12. Hosono K, Noda S, Shimizu A, Nakanishi N, Ohtsubo M, Shimizu N, et al. YPEL5 protein of the YPEL gene family is involved in the cell cycle progression by interacting with two distinct proteins RanBPM and RanBP10. Genomics. 2010;96(2):102–11. doi: 10.1016/j.ygeno.2010.05.003. PMID: 20580816

13. Ciuladaite Z, Kasnauskiene J, Cimbalistiene L, Preiksaitiene E, Patsalis P, Kucinskas V. Mental retardation and autism associated with recurrent 16p11.2 microdeletion: incomplete penetrance and variable expressivity. JOURNAL OF APPLIED GENETICS. 2011;52(4):443–9. doi: 10.1007/s13353-011-0063-z. PMID: 21931978

14. Sebat J, Lakshmi B, Malhotra D, Troge J, Lese-Martin C, Walsh T, et al. Strong association of de novo copy number mutations with autism. SCIENCE. 2007;316(5823):445–9. doi: 10.1126/science.1138659. PMID: 17363630

15. Kumar R, KaraMohamed S, Sudi J, Conrad D, Brune C, Badner J, et al. Recurrent 16p11.2 microdeletions in autism. HUMAN MOLECULAR GENETICS. 2008;17(4):628–38. doi: 10.1093/hmg/ddm376. PMID: 18156158

16. Maillard A, Ruef A, Pizzagalli F, Migliavacca E, Hippolyte L, Adaszewski S, et al. The 16p11.2 locus modulates brain structures common to autism, schizophrenia and obesity. MOLECULAR PSYCHIATRY. 2015;20(1):140–7. doi: 10.1038/mp.2014.145. PMID: 25421402

17. McCarthy S, Makarov V, Kirov G, Addington A, McClellan J, Yoon S, et al. Microduplications of 16p11.2 are associated with schizophrenia. NATURE GENETICS. 2009;41(11):1223–U85. doi: 10.1038/ng.474. PMID: 19855392

18. Roll P, Sanlaville D, Cillario J, Labalme A, Bruneau N, Massacrier A, et al. Infantile Convulsions with Paroxysmal Dyskinesia (ICCA Syndrome) and Copy Number Variation at Human Chromosome 16p11. PLOS ONE. 2010;5(10). doi: 10.1371/journal.pone.0013750. PMID: 21060786

19. Blaker-Lee A, Gupta S, McCammon JM, De Rienzo G, Sive H. Zebrafish homologs of genes within 16p11.2, a genomic region associated with brain disorders, are active during brain development, and include two deletion dosage sensor genes. Disease Models & Mechanisms. 2012;5(6):834–51. doi: 10.1242/dmm.009944. PMID: 22566537

20. Blanco-Sanchez B, Clement A, Stednitz SJ, Kyle J, Peirce JL, McFadden M, et al. yippee like 3 (ypel3)is a novel gene required for myelinating and perineurial glia development. Plos Genetics. 2020;16(6). doi: 10.1371/journal.pgen.1008841. PMID: 32544203

21. Kim JH, Singh M, Pan G, Lopez A, Zito N, Bosse B, et al. Frameshift mutations of YPEL3 alter the sensory circuit function in Drosophila. Disease Models & Mechanisms. 2020;13(6). doi: 10.1242/dmm.042390. PMID: 32461240

22. Zatloukal K, Stumptner C, Fuchsbichler A, Heid H, Schnoelzer M, Kenner L, et al. p62 is a common component of cytoplasmic inclusions in protein aggregation diseases. AMERICAN JOURNAL OF PATHOLOGY. 2002;160(1):255–63. PMID: 11786419

23. Foster A, Rea S. The role of sequestosome 1/p62 protein in amyotrophic lateral sclerosis and frontotemporal dementia pathogenesis. NEURAL REGENERATION RESEARCH. 2020;15(12):2186–94. doi: 10.4103/1673-5374.284977. PMID: 32594029

24. Muto V, Flex E, Kupchinsky Z, Primiano G, Galehdari H, Dehghani M, et al. Biallelic SQSTM1 mutations in early-onset, variably progressive neurodegeneration. EUROPEAN JOURNAL OF HUMAN GENETICS. 2019;27:311–2. PMID: 29959261

25. Haack T, Ignatius E, Calvo-Garrido J, Iuso A, Isohanni P, Maffezzini C, et al. Absence of the Autophagy Adaptor SQSTM1/p62 Causes Childhood-Onset Neurodegeneration with Ataxia, Dystonia, and Gaze Palsy. AMERICAN JOURNAL OF HUMAN GENETICS. 2016;99(3):735–43. doi: 10.1016/j.ajhg.2016.06.026. PMID: 27545679

26. Lamark T, Svenning S, Johansen T. Regulation of selective autophagy: the p62/SQSTM1 paradigm. SIGNALLING MECHANISMS IN AUTOPHAGY2017. p. 609–24. doi: 10.1042/EBC20170035.PMID: 29233872

27. Katsuragi Y, Ichimura Y, Komatsu M. p62/SQSTM1 functions as a signaling hub and an autophagy adaptor. FEBS JOURNAL. 2015;282(24):4672–8. doi: 10.1111/febs.13540. PMID: 26432171

28. Damme M, Suntio T, Saftig P, Eskelinen E. Autophagy in neuronal cells: general principles and physiological and pathological functions. ACTA NEUROPATHOLOGICA. 2015;129(3):337–62. doi: 10.1007/s00401-014-1361-4. PMID: 25367385

29. Oswald M, Garnham N, Sweeney S, Landgraf M. Regulation of neuronal development and function by ROS. FEBS LETTERS. 2018;592(5):679–91. doi: 10.1002/1873-3468.12972. PMID: 29323696

30. Odell S, Clark D, Zito N, Jain R, Gong H, Warnock K, et al. Internal state affects local neuron function in an early sensory processing center to shape olfactory behavior in Drosophila larvae. SCIENTIFIC REPORTS. 2022;12(1). doi: 10.1038/s41598-022-20147-1. PMID: 36131078

31. Guruharsha K, Rual J, Zhai B, Mintseris J, Vaidya P, Vaidya N, et al. A Protein Complex Network of Drosophila melanogaster. CELL. 2011;147(3):690–703. doi: 10.1016/j.cell.2011.08.047. PMID: 22036573

32. Turan G, Olgun Ç, Ayten H, Toker P, Ashyralyyev A, Savas B, et al. Dynamic proximity interaction profiling suggests that YPEL2 is involved in cellular stress surveillance. PROTEIN SCIENCE. 2024;33(2). doi: 10.1002/pro.4859. PMID: 38145972

33. Nezis I, Simonsen A, Sagona A, Finley K, Gaumer S, Contamine D, et al. Ref(2) P, the Drosophila melanogaster homologue of mammalian p62, is required for the formation of protein aggregates in adult brain. JOURNAL OF CELL BIOLOGY. 2008;180(6):1065–71. PMID: 18347073

34. Grueber W, Ye B, Yang C, Younger S, Borden K, Jan L, et al. Projections of Drosophila multidendritic neurons in the central nervous system:: links with peripheral dendrite morphology. DEVELOPMENT. 2007;134(1):55–64. doi: 10.1242/dev.02666. PMID: 17164414

35. Xie Z, Klionsky D. Autophagosome formation: Core machinery and adaptations. NATURE CELL BIOLOGY. 2007;9(10):1102–9. doi: 10.1038/ncb1007-1102. PMID: 17909521

36. Sanyal S. Genomic mapping and expression patterns of C380, OK6 and D42 enhancer trap lines in the larval nervous system of Drosophila. GENE EXPRESSION PATTERNS. 2009;9(5):371–80. doi: 10.1016/j.gep.2009.01.002. PMID: 19602393

37. Kinghorn K, Grönke S, Castillo-Quan J, Woodling N, Li L, Sirka E, et al. A Drosophila Model of Neuronopathic Gaucher Disease Demonstrates Lysosomal-Autophagic Defects and Altered mTOR Signalling and Is Functionally Rescued by Rapamycin. JOURNAL OF NEUROSCIENCE. 2016;36(46):11654–70. doi: 10.1523/JNEUROSCI.4527-15.2016. PMID: 27852774

38. Bell K, Al-Mubarak B, Martel M, McKay S, Wheelan N, Hasel P, et al. Neuronal development is promoted by weakened intrinsic antioxidant defences due to epigenetic repression of Nrf2. NATURE COMMUNICATIONS. 2015;6. doi: 10.1038/ncomms8066. PMID: 25967870

39. Albrecht S, Barata A, Grosshans J, Teleman A, Dick T. In Vivo Mapping of Hydrogen Peroxide and Oxidized Glutathione Reveals Chemical and Regional Specificity of Redox Homeostasis. CELL METABOLISM. 2011;14(6):819–29. doi: 10.1016/j.cmet.2011.10.010. PMID: 22100409

40. Zhang J, Wen X, Ren X, Li Y, Tang X, Wang Y, et al. YPEL3 suppresses epithelial-mesenchymal transition and metastasis of nasopharyngeal carcinoma cells through the Wnt/β-catenin signaling pathway. JOURNAL OF EXPERIMENTAL & CLINICAL CANCER RESEARCH. 2016;35. doi: 10.1186/s13046-016-0384-1. PMID: 27400785

41. Bradley C, Peineau S, Taghibiglou C, Nicolas C, Whitcomb D, Bortolotto Z, et al. A pivotal role of GSK-3 in synaptic plasticity. FRONTIERS IN MOLECULAR NEUROSCIENCE. 2012;5. doi: 10.3389/fnmol.2012.00013. PMID: 22363262

42. Franco B, Bogdanik L, Bobinnec Y, Debec A, Bockaert J, Parmentier M, et al. Shaggy, the homolog of glycogen synthase kinase 3, controls neuromuscular junction growth in Drosophila. JOURNAL OF NEUROSCIENCE. 2004;24(29):6573–7. doi: 10.1523/JNEUROSCI.1580-04.2004. PMID: 15269269

43. Babu J, Seibenhener M, Peng J, Strom A, Kemppainen R, Cox N, et al. Genetic inactivation of p62 leads to accumulation of hyperphosphorylated tau and neurodegeneration. JOURNAL OF NEUROCHEMISTRY. 2008;106(1):107–20. doi: 10.1111/j.1471-4159.2008.05340.x. PMID: 18346206

44. Shen W, Ganetzky B. Autophagy promotes synapse development in Drosophila. JOURNAL OF CELL BIOLOGY. 2009;187(1):71–9. doi: 10.1083/jcb.200907109. PMID: 19786572

45. Gottemukkala K, Chrustowicz J, Sherpa D, Sepic S, Vu D, Karayel O, et al. Non-canonical substrate recognition by the human WDR26-CTLH E3 ligase regulates prodrug metabolism. MOLECULAR CELL. 2024;84(10). doi: 10.1016/j.molcel.2024.04.014. PMID: 38759627

46. Ramsey C, Glass C, Montgomery M, Lindl K, Ritson G, Chia L, et al. Expression of Nrf2 in neurodegenerative diseases. JOURNAL OF NEUROPATHOLOGY AND EXPERIMENTAL NEUROLOGY. 2007;66(1):75–85. PMID: 17204939

47. Zhao F, Wu T, Lau A, Jiang T, Huang Z, Wang X, et al. Nrf2 promotes neuronal cell differentiation. FREE RADICAL BIOLOGY AND MEDICINE. 2009;47(6):867–79. doi: 10.1016/j.freeradbiomed.2009.06.029. PMID: 19573594

48. Xiong W, Garfinkel A, Li Y, Benowitz L, Cepko C. NRF2 promotes neuronal survival in neurodegeneration and acute nerve damage. JOURNAL OF CLINICAL INVESTIGATION. 2015;125(4):1433–45. doi: 10.1172/JCI79735. PMID: 25798616

49. Branca C, Ferreira E, Nguyen T, Doyle K, Caccamo A, Oddo S. Genetic reduction of Nrf2 exacerbates cognitive deficits in a mouse model of Alzheimer’s disease. HUMAN MOLECULAR GENETICS. 2017;26(24):4823–35. doi: 10.1093/hmg/ddx361. PMID:

50. Ren P, Chen J, Li B, Zhang M, Yang B, Guo X, et al. Nrf2 Ablation Promotes Alzheimer’s Disease-Like Pathology in APP/PS1 Transgenic Mice: The Role of Neuroinflammation and Oxidative Stress. OXIDATIVE MEDICINE AND CELLULAR LONGEVITY. 2020;2020. doi: 10.1155/2020/3050971.PMID: 29036636

51. Chen L, Yue Z, Liu Z, Liu H, Zhang J, Zhang F, et al. The impact of Nrf2 knockout on the neuroprotective effects of dexmedetomidine in a mice model of cognitive impairment. BEHAVIOURAL BRAIN RESEARCH. 2024;469. doi: 10.1016/j.bbr.2024.115006.PMID: 38692357

52. Oswald M, Brooks P, Zwart M, Mukherjee A, West R, Giachello C, et al. Reactive oxygen species regulate activity-dependent neuronal plasticity in Drosophila. ELIFE. 2018;7. doi: 10.7554/eLife.39393.PMID: 30540251

53. Doser R, Knight K, Deihl E, Hoerndli F. Activity-dependent mitochondrial ROS signaling regulates recruitment of glutamate receptors to synapses. ELIFE. 2024;13. doi: 10.7554/eLife.92376. PMID: 38483244

54. Levings D, Pathak S, Yang Y, Slattery M. Limited expression of Nrf2 in neurons across the central nervous system. REDOX BIOLOGY. 2023;65. doi: 10.1016/j.redox.2023.102830. PMID: 37544245

55. Stavrovskaya I, Morin B, Madamba S, Alexander C, Romano A, Alam S, et al. Mitochondrial ROS modulate presynaptic plasticity in the drosophila neuromuscular junction. REDOX BIOLOGY. 2025;79. doi: 10.1016/j.redox.2024.103474. PMID: 39721493

56. Tuttle R, Simon M, Hitch D, Maiorano J, Hellan M, Ouellette J, et al. Senescence-Associated Gene YPEL3 Is Downregulated in Human Colon Tumors. ANNALS OF SURGICAL ONCOLOGY. 2011;18(6):1791–6. doi: 10.1245/s10434-011-1558-x. PMID: 21267786

57. Wang L, Zhang Z, Zhou X, Wu J, Hong Z. Comprehensive analysis of the expression and prognosis of YPEL family members in clear cell renal cell cancer. ONCOLOGY REPORTS. 2022;48(1). doi: 10.3892/or.2022.8345. PMID: 35674183

58. Lei C, Zhao B, Liu L, Zeng X, Yu Z, Wang X. Expression and clinical significance of p62 protein in colon cancer. MEDICINE. 2020;99(3). doi: 10.1097/MD.0000000000018791. PMID: 32011477

59. Philipson E, Engström C, Naredi P, Fagman J. High expression of p62/SQSTM1 predicts shorter survival for patients with pancreatic cancer. BMC CANCER. 2022;22(1). doi: 10.1186/s12885-022-09468-6. PMID: 35354432

60. Tang J, Li Y, Xia S, Li J, Yang Q, Ding K, et al. Sequestosome 1/p62: A multitasker in the regulation of malignant tumor aggression (Review). INTERNATIONAL JOURNAL OF ONCOLOGY. 2021;59(4). doi: 10.3892/ijo.2021.5257. PMID: 34414460

61. Sporn M, Liby K. NRF2 and cancer: the good, the bad and the importance of context. NATURE REVIEWS CANCER. 2012;12(8):564–71. doi: 10.1038/nrc3278. PMID: 22810811

62. Pouremamali F, Pouremamali A, Dadashpour M, Soozangar N, Jeddi F. An update of Nrf2 activators and inhibitors in cancer prevention/promotion. CELL COMMUNICATION AND SIGNALING. 2022;20(1). doi: 10.1186/s12964-022-00906-3. PMID: 35773670

63. Robledinos-Antón N, Fernández-Ginés R, Manda G, Cuadrado A. Activators and Inhibitors of NRF2: A Review of Their Potential for Clinical Development. OXIDATIVE MEDICINE AND CELLULAR LONGEVITY. 2019;2019. doi: 10.1155/2019/9372182. PMID: 31396308

64. Kim J, Wang X, Coolon R, Ye B. Dscam Expression Levels Determine Presynaptic Arbor Sizes in Drosophila Sensory Neurons. NEURON. 2013;78(5):827–38. doi: 10.1016/j.neuron.2013.05.020. PMID: 23764288

65. Kim S, Quagraine Y, Singh M, Kim J. Rab11 suppresses neuronal stress signaling by localizing dual leucine zipper kinase to axon terminals for protein turnover. ELIFE. 2024;13. doi: 10.7554/eLife.96592. PMID: 39475475

66. Castells-Nobau A, Nijhof B, Eidhof I, Wolf L, Scheffer-de Gooyert J, Monedero I, et al. Two Algorithms for High-throughput and Multi-parametric Quantification of Drosophila Neuromuscular Junction Morphology. JOVE-JOURNAL OF VISUALIZED EXPERIMENTS. 2017;(123). doi: 10.3791/55395. PMID: 28518121

67. Schindelin J, Arganda-Carreras I, Frise E, Kaynig V, Longair M, Pietzsch T, et al. Fiji: an open-source platform for biological-image analysis. NATURE METHODS. 2012;9(7):676–82. doi: 10.1038/NMETH.2019. PMID: 22743772

68. Gamarra M, Blanco-Urrejola M, Batista A, Imaz J, Baleriola J. Object-Based Analyses in FIJI/ImageJ to Measure Local RNA Translation Sites in Neurites in Response to Aβ1-42 Oligomers. FRONTIERS IN NEUROSCIENCE. 2020;14. doi: 10.3389/fnins.2020.00547. PMID: 32581689

